# Multiciliated cells adapt the mechanochemical Piezo1-Erk1/2-Yap1 cell proliferation axis to fine-tune centriole number

**DOI:** 10.1101/2025.01.21.634139

**Authors:** Vani Narayanan, Venkatramanan G Rao, Angelo Arrigo, Saurabh S Kulkarni

## Abstract

Multiciliated cells (MCCs) are specialized epithelial cells that undergo massive amplification of centrioles, constructing several motile cilia to propel fluid flow. The abundance of cilia is critical for efficient fluid flow, yet how MCCs regulate centriole/cilia numbers remains a major knowledge gap. We have shown that mechanical tension plays a central role in regulating apical area and centriole number in MCCs. Here, we demonstrate that centriole amplification is controlled by a mechanochemical pathway essential for cell proliferation in cycling cells. Specifically, MCCs under tension use Piezo1-mediated calcium signaling to drive Erk½ phosphorylation via PKC and subsequent Yap1 activation. Remarkably, MCCs use this pathway to activate a cilia-specific transcription program, influencing the expression of Foxj1, a master regulator of motile ciliogenesis. Our work is the first to identify a novel function for an important mechanochemical pathway in centriole amplification in MCCs, offering new insights into ciliopathies and cancer, where aberrant centriole numbers are implicated.

**Teaser:** This study demonstrates that multiciliated cells utilize the mechanochemical Piezo1-Erk1/2-Yap1 cell proliferation axis to activate the cilia-specific transcriptional factor Foxj1 and amplify centrioles in a tension-dependent manner.

## INTRODUCTION

The differentiation of progenitor cells into specialized cells involves significant remodeling of their transcriptional landscape and a definitive exit from the cell cycle. Among the most striking instances of this transformation are multiciliated cells (MCCs). These cells notably redirect their developmental trajectory to avoid mitotic commitment and instead undergo massive centriole amplification, enabling them to build multiple motile cilia. These cilia coordinate hydrodynamic fluid movement, which is crucial for processes like mucociliary clearance. Mutations that affect the number of cilia/centrioles are linked to significant health issues, including progressive lung disorders (*1, 2*), infertility (*3, 4*), and hydrocephalus (*4, 5*), posing the key question: how do MCCs amplify the right number of centrioles?

Cells and tissues undergoing differentiation and morphogenesis are frequently stretched, compressed, or subjected to mechanical forces. Cells respond by translating mechanical cues into chemical signals using mechanosensitive proteins to ensure proper tissue formation and function (*6, 7*). A classic example is when epithelial cells are subjected to mechanical tension; they undergo centriole duplication and cell division to maintain optimal cell number and barrier function (*8–10*). MCCs, albeit epithelial, are post-mitotic and thus do not undergo cell division when subjected to mechanical tension. Instead, MCCs increase in apical area and undergo centriole amplification (*11–13*). How MCCs exhibit a different response may depend on the mechanochemical signaling these cells initiate.

In epithelial cells, mechanical tension activates a stretch-activated channel, Piezo1, elevating intracellular Ca^2+^ levels. This influx of Ca^2+^ leads to the activation of a MAPK, ERK ½, thereby driving cells into mitosis (*9*). While studies show an interplay between MAPK and Hippo signaling cascades (*14*) to promote cellular proliferation in response to mechanical strain, Piezo1 activation also increases the activity of a co-transcriptional activator, YAP, and, hence, promotes cell proliferation and tumorigenesis (*15, 16*). Recent findings also highlight the role of Piezo1 in coordinating with MAPK and YAP signaling to enhance tumor growth in hepatocellular carcinoma (*17*).

We recently showed that MCCs use Piezo1 in the multiciliated epithelium of the *Xenopus* embryonic epidermis to control centriole amplification in response to stretch. In this study, we demonstrate that, though MCCs do not undergo cell division, they use the same Piezo1-MAPK-YAP mechanochemical axis to respond to mechanical strain. They divert from cycling epithelial cells by activating the cilia-specific gene transcription program, specifically, the expression of Foxj1, a master regulator of motile ciliogenesis, to control centriole and cilia number in MCCs. Our findings uncover a novel function for an important mechanochemical pathway, underscoring a unique aspect of MCC morphogenesis and providing crucial insights into the molecular basis of conditions such as primary ciliary dyskinesia, hydrocephalus, and infertility.

## RESULTS

### Piezo1 regulates centriole numbers in a cell-autonomous manner

Global depletion of Piezo1 leads to a reduction in centriole number and an increase in the apical area in a tension-dependent manner, resulting in a confounding effect (*11*). While the apical area of MCCs is determined by the interplay between cell intrinsic and extrinsic forces, the centriole amplification program is specific to MCCs (*11, 18–20*). To test the role of Piezo1 in centriole amplification, we generated a mosaic knockdown (KD) of Piezo1 by injecting Piezo1 MO (*11*) with H2B-RFP as a tracer in one of the four cells at the 4-cell stage embryo (Figure 1A). Embryos were then analyzed for four possible outcomes. First, Piezo1 is not KD in MCCs and surrounding non-MCCs. Second, Piezo1 is KD in an MCC but not the surrounding non-MCCs. Third, Piezo1 was present in an MCC but was KD in surrounding non-MCCs. Fourth, Piezo1 was KD in MCC and surrounding non-MCCs (Figure 1B). Using this strategy, we show that centriole number and density (number of centrioles per unit area of MCC) are affected when Piezo1 is depleted in MCCs (Scenario #2 and #4) but not when Piezo1 is depleted in non-MCCs only (Scenario #3) (Fig. 1, C and D, and fig. S1A). The reduction in centriole density in #4 is due to both a decrease in centriole number and an increase in the apical area, similar to the global depletion of Piezo1. Whereas, when Piezo1 is depleted in MCCs alone (#2), it only affects the number of centrioles but not the apical area of MCCs, leading to a decrease in centriole density. In contrast, the apical surface area is affected only when Piezo1 is lost in both MCCs and non-MCCs (Scenario #4). Thus, our results demonstrate that Piezo1 regulates centriole number, but not the apical area, in an MCC cell-autonomous manner.

**Figure 1.**
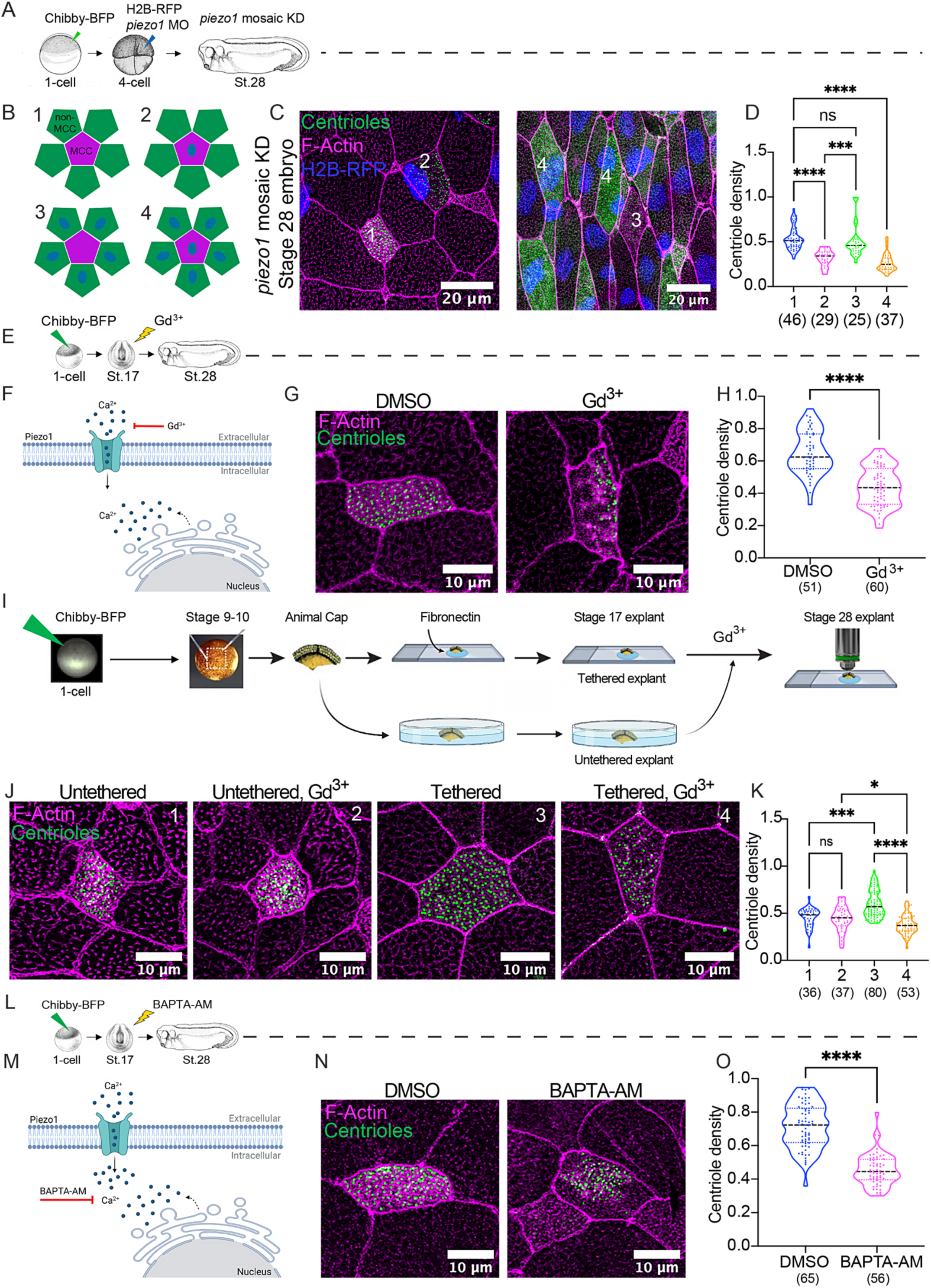
Role of Piezo1 and Calcium in centriole amplification in *Xenopus* MCCs. A. Schematic representing the experimental design for mosaic knockdown of Piezo1. Embryos were injected with a centriole marker (Chibby-BFP, 100 pg, green) at 1-cell stage, followed by injection with piezo1 morpholino and a nuclear tracer (H2B-RFP, 100 pg, blue) to trace the knockdown at 1 in 4 cell stage, and then allowed to develop until stage 28. B. Schematic representing the four cases yielded by the mosaic knockdown approach yielded four cases. (1) control, (2) Only MCCs are affected by the knockdown, (3) Only non-MCC’s are depleted of Piezo1 and MCCs are unaffected, and (4) both, MCCs and non-MCCs are depleted of Piezo1. C. Representative immunofluorescence (IF) images of piezo1 mosaic KD stage 28 embryos expressing Chibby-BFP (centrioles, green), H2B-RFP (nuclear tracer, blue), and co-stained with Phalloidin (Magenta). Left: IF image depicts cases 1 and 2, Right: IF image depicts cases 3 and 4. Scale bar: 20um. D. Plot shows centriole density per MCC (centriole number/apical area) for all four cases. Kruskal-Wallis test followed by multiple comparisons was performed for statistical analysis between groups (p>0.05 (ns), ***p<0.001, and ****p<0.0001). Data shown is from three independent experiments; number of MCCs from at least 5 embryos per trial is indicated in parentheses. E. Schematic of the experimental design for gadolinium chloride treatment. Embryos were injected with Chibby-BFP (green) at 1-cell stage, treated with 30uM of Gadolinium Chloride at stage 17, and collected at stage 28 for immunostaining and visualization. F. Schematic represents the mechanism by which gadolinium chloride blocks Piezo1- mediated calcium influx. G. Representative IF images of stage 28 MCCs that are either untreated (Control) or treated with gadolinium chloride (Gd^3+^), expressing Chibby-BFP (centrioles, green) and are co-stained with Phalloidin (Magenta). Scale bar: 10um. H. Plot depicts comparison of centriole density per MCC between control and Gd^3+^-treated stage 28 embryos. Mann-Whitney non-parametric t-test was used for statistical analysis (****p<0.0001). Data shown is from three independent experiments; number of MCCs from at least 5 embryos per group per trial is indicated in parentheses. I. Schematic representing the experimental design for animal explant extraction and drug treatment with 30uM of Gadolinium Chloride (Gd^3+^) prior to staining and visualization. Animal explants were surgically extracted at stage 9-10 from embryos that were injected with Chibby-BFP (green) at 1-cell stage, followed by culture in either fibronectin-coated slides (tethered) or an uncoated dish with media (untethered). The explants were then exposed to Gd^3+^ at stage 17 and allowed to develop until stage 28. The explants were stage-matched with unmanipulated sibling embryos. J. Representative IF images of stage 28 MCCs in animal explants expressing Chibby-BFP (centrioles, green) and co-stained with Phalloidin (Magenta). The tethered explant treated with Gd^3+^ (Tethered, Gd^3+^) is compared against the untreated tethered control (Tethered) and the untethered groups (Untethered and Untethered, Gd^3+^). Scale bar is 10um. K. Plot shows comparison of the centriole density per MCC between the different groups in (J). Kruskal-Wallis test followed by multiple comparisons was performed for statistical analysis between groups (p>0.05 (ns), ***p<0.001, and ****p<0.001). Data shown is from three independent experiments; the number of MCCs from at least 5 animal explants per group per experiment is indicated in parentheses below each group. L. Schematic of the experimental design for BAPTA-AM treatment. Embryos were injected with Chibby-BFP (green) at 1-cell stage, treated with 100uM of BAPTA-AM at stage 17, and collected at stage 28 for immunostaining and visualization. M. Schematic represents the mechanism by which the intra-cellular calcium chelator, BAPTA-AM blocks intra-cellular calcium inside the cell. N. Representative IF images of stage 28 MCCs that are either untreated (Control) or treated with BAPTA-AM (BAPTA-AM), expressing Chibby-BFP (centrioles, green) and are co-stained with Phalloidin (Magenta). Scale bar: 10um.

Plot depicts a comparison of centriole density per MCC for control and BAPTA-AM-treated stage 28 embryos. Mann-Whitney non-parametric t-test was used for statistical analysis (****p<0.0001). Data shown is from three independent experiments; the number of MCCs from at least 5 embryos per group per experiment is indicated in parentheses below each group.

### Piezo1-mediated calcium influx is essential for centriole amplification in MCCs

Mounting evidence suggests Piezo1 is a mechanosensitive calcium channel (*21–23*). Therefore, we hypothesized that Piezo1-mediated calcium influx contributes to centriole amplification in MCCs. To test this hypothesis, we treated *Xenopus* embryos with gadolinium, a potent inhibitor (competitor) of calcium channels (*24, 25*). While gadolinium is not a specific inhibitor of Piezo1-mediated calcium influx, gadolinium treatment in whole embryos significantly reduced centriole number and density in MCCs, similar to Piezo1 knockdown, knockout, and GSMTx4 treatments (*11*) (Fig. 1, E to H, fig. S1B). Next, we used tethered and untethered animal cap explants to test whether the effect of gadolinium on centriole amplification is tension-dependent. Indeed, gadolinium treatment blocked the amplification of centrioles only in tethered explants (Fig. 1, I to K, fig. S1C), underscoring the potential role of calcium influx in regulating tension-dependent centriole amplification in MCCs.

To address the possibility of off-target effects of gadolinium, we directly probed the role of calcium in centriole amplification. Removing extracellular calcium using divalent-free media (DMSO in Ca^2+^ and Mg ^2+^ - free media) slightly but significantly reduced the centriole number and apical area, keeping centriole density the same as controls (fig. S1, D, and E). This result suggests a role for extracellular calcium in apical area and centriole amplification. However, intracellular calcium may compensate for the lack of calcium influx from the medium, affecting the significance of our results. While EGTA and BAPTA are widely used as calcium-chelating reagents, BAPTA seldom deprotonates in physiological pH, leading to a higher complexation rate with calcium than EGTA (*26, 27*). Therefore, to test the hypothesis, we chelated intracellular calcium using a cell-permeant form of BAPTA, BAPTA-AM, in the presence and absence of extracellular calcium. Both BAPTA-AM alone (BAPTA-AM) and BAPTA-AM in divalent-free media (B-AM in Ca^2+^ and Mg ^2+^ - free media) significantly reduced centriole number and centriole density, supporting the hypothesis that calcium is essential for centriole amplification in MCCs (Fig. 1, L to O, fig. S1, D to F). We note that sometimes centrioles are clumped in these experiments, possibly due to defects in apical expansion or enrichment of F-actin at the apical surface, making it harder to count centrioles accurately. Therefore, we use IMARIS for segmentation and 3D rendering to visualize and count centrioles more accurately. Given the extensive use of BAPTA and its derivatives in calcium signaling research, it is important to note that these molecules manifest off-target effects that affect cytoskeleton dynamics independent of the calcium chelating property (*28, 29*). Therefore, we sought to investigate the role of calcium signaling in centriole number regulation further by examining the downstream effectors of calcium.

### Piezo1-mediated calcium influx activates the PKC-Erk½ pathway to control centriole number

The mitogen-activated protein kinase (Erk, Jnk, and p38) pathway is an important bridge in the switch from extracellular mechanical signals to intracellular biochemical responses. Specifically, mechanical stretch is shown to activate the Erk½ pathway via Piezo1-mediated calcium influx to promote epithelial cell proliferation (*9*). Since MCCs are post-mitotic, we reasoned that MCCs might adapt the Piezo1-Erk½ cell proliferation pathway to amplify centrioles in response to mechanical forces. To test this hypothesis, we stimulated Piezo1 via a channel-specific chemical agonist, Yoda1 (*30, 31*) (fig. S2A). As expected, Yoda1 increased phosphorylated Erk½ relative to total Erk½ levels in stage 28 embryos (fig. S2, B, and D). Prior studies in other model systems have shown Erk½ activation to correlate with its nuclear localization (*6, 32–34*). Indeed, Piezo1 activation via Yoda1 increased nuclear localization of phosphorylated Erk½ in MCCs (fig. S2, C, and E).

**Figure 2.**
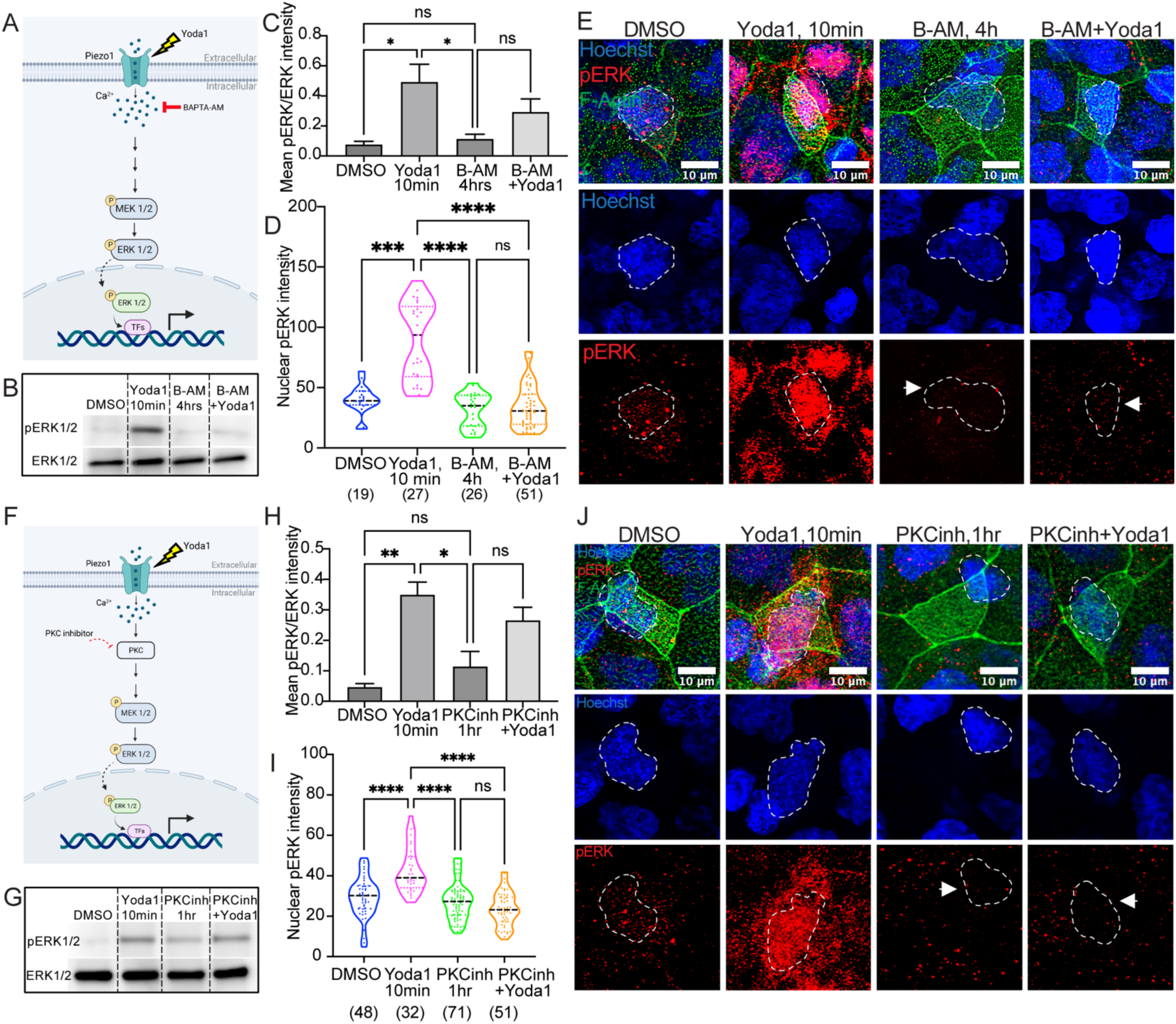
Piezo1-mediated calcium influx activates the PKC-Erk½ pathway in *Xenopus* MCCs. A. Schematic represents the mechanism of Piezo1 activation via Yoda1 and subsequent activation of the ERK ½ pathway indicated by the nuclear translocation of phosphorylated ERK ½ to the nucleus. The diagram also depicts the mechanism of intracellular calcium chelation via BAPTA-AM. B. Western analysis showing the effect of BAPTA-AM (100 uM) and Yoda1 (25 uM) treatment on ERK ½ phosphorylation. Embryos were either treated with BAPTA-AM alone for 4 hours (B-AM) or pre-treated with BAPTA-AM for 4 hours followed by BAPTA-AM and Yoda1 treatment for 10 minutes (B-AM + Yoda1). The treated embryos were collected at stage 28 and compared for phospho- ERK ½ and total ERK ½ expression against stage 28 embryos treated with Yoda1 alone for 10 minutes (Yoda1) and the DMSO-treated vehicle control (DMSO). Representative data shown is from one experiment out of three independent experiments. C. Plot shows mean intensity of phospho-ERK ½ versus total ERK ½ across the different treatment groups in (B). One-Way ANOVA test followed by multiple comparisons was performed for statistical analysis between groups (p>0.05 (ns), ***p<0.001, and ****p<0.0001). Data shown is from three independent experiments; 30 embryos were used per group per trial. D. Plot shows intensity of nuclear phospho-ERK ½ per MCC in stage 28 embryos that were either DMSO-treated (DMSO), treated with Yoda1 for 10 minutes (Yoda1, 10min), treated with BAPTA-AM alone for 4 hours (B-AM, 4h), or pre-treated with BAPTA-AM for 4 hours followed by BAPTA-AM and Yoda1 for 10 minutes (B-AM+Yoda1). Kruskal-Wallis test followed by multiple comparisons was performed for statistical analysis between groups (p>0.05 (ns), ***p<0.001, and ****p<0.0001). Data shown is from three independent experiments. The number of MCCs used for each group from every trial is indicated in parentheses. At least 5 embryos were used per group in every experiment. E. Representative IF images show nuclear expression of phospho-ERK ½ (pERK, red) in MCCs across the different treatment groups in (D). Embryos were co-stained with Phalloidin (F-Actin, green) and Hoechst (nuclei, blue). Scale bar is 10 um. F. Schematic represents the mechanism of Piezo1 activation of ERK ½ pathway via Yoda1 and protein kinase C (PKC)-mediated pathway inhibition via the PKC inhibitor (Bisindolylmaleimide II or BIM-II). G. Western analysis showing the effect of PKC inhibitor (25 uM) and Yoda1 (25 uM) treatment on ERK ½ phosphorylation. Embryos were either treated with PKC inhibitor alone for 1 hour (PKCinh) or pre-treated with PKC inhibitor for 1 hour followed by PKC inhibitor and Yoda1 treatment for 10 minutes (PKCinh + Yoda1). The treated embryos were collected at stage 28 and compared for phospho-ERK ½ and total ERK ½ expression against stage 28 embryos treated with Yoda1 alone for 10 minutes (Yoda1) and the DMSO-treated vehicle control (DMSO). Representative data shown is from one experiment out of three independent experiments. H. Plot shows mean intensity of phospho-ERK ½ versus total ERK ½ across the different treatment groups in (G). One-Way ANOVA test followed by multiple comparisons was performed for statistical analysis between groups (p>0.05 (ns), **p<0.01, and ****p<0.0001). Data shown is from three independent experiments; 30 embryos were used per group per trial. I. Plot shows intensity of nuclear phospho-ERK ½ per MCC from stage 28 embryos that were either vehicle-treated (DMSO), treated with Yoda1 for 10 minutes (Yoda1, 10min), treated with PKC inhibitor alone for 1 hour (PKCinh, 1hr), or pre-treated with PKC inhibitor for 1 hour followed by PKC inhibitor and Yoda1 for 10 minutes (PKCinh+Yoda1). Kruskal-Wallis test followed by multiple comparisons was performed for statistical analysis between groups (p>0.05 (ns), **p<0.01, and ****p<0.0001). Data shown is from three independent experiments; the number of MCCs from at least 5 embryos per trial is indicated in parentheses below each group. Representative IF images show nuclear expression of phospho-ERK ½ (pERK, red) in MCCs across the different treatment groups in (I). Embryos were co-stained with Phalloidin (F-Actin, green) and Hoechst (nuclei, blue). Scale bar is 10 um.

Next, we determined whether Piezo1-mediated calcium influx and its downstream effectors were required for phosphorylation and subsequent nuclear entry of Erk½ in MCCs. Piezo1 stimulation in the presence of calcium chelator, BAPTA-AM (Fig. 2, A to E) or divalent-free media (DVF) to remove extracellular calcium (fig. S2, F to J), or PKC inhibition via Bisindolylmaleimide II (BIM-II) (Fig. 2, F to J), remarkably reduced activation of Erk½ in MCCs, suggesting an essential role of Piezo1-mediated calcium influx in triggering the Erk½ cascade in MCCs. It is important to note that the western blot experiments were performed on whole embryos consisting of different types of cells, yielding a global effect on Erk ½ phosphorylation. Hence, our results were orthogonally validated with the help of spatially resolved immunofluorescence staining that was more specific to MCCs (Fig. 2, D and E, I and J, fig. S2, H, and J).

Next, we examined if Erk½ is essential for centriole amplification in MCCs, as it is activated (by phosphorylation) under mechanical load in *Xenopus* embryonic epidermis (*6*). Specifically, Erk2, compared to Erk1, was shown to be mainly involved in differentiation and organismal development (*35*). Depletion of Erk2 via morpholino-based knockdown partly affected gastrulation and the anterior-posterior elongation of the embryos (fig. S3, A and B). However, most embryos could reach stage 28 when we assess the MCC development. To our surprise, Erk2- depleted embryos could differentiate MCCs, but depletion of Erk2 blocked centriole amplification (as indicated by fewer centrioles), reduced apical area, and reduced centriole density (Fig. 3, A to C, and fig. S3, C and D). Next, we verified *erk2* MO specificity using a mosaic rescue experiment. Specifically, we injected *erk2* MO at the 1-cell stage to deplete Erk2 from the whole embryo and injected WT mCherry-*ERK2* in 1 of the 4 cells at the 4- cell stage to rescue centriole number in MCCs that receive mCherry-*ERK2* (Fig. 3D). Indeed, these defects were partially rescued with human WT mCherry-*ERK2* (Fig. 3, E and F). To avoid early developmental defects caused by Erk2 depletion, we decided to inhibit the upstream regulators of Erk½ cascade, PKC (via BIM-II), or Mek½ (via PD0325901) at stage 17, when embryos have finished gastrulation and have begun to differentiate the MCCs. We observed interesting differences in their effects on apical expansion, with Mek½ inhibition leading to a modest increase and PKC inhibition leading to a significant decrease in the apical areas, similar to Erk2 knockdown (fig. S3, F and H). We do not understand the reason behind these differences, but multiple factors could play a role. For example, the timing of Erk½ inhibition is much later compared to *erk2* MO, Mek½ affects both Erk½ and not just Erk2 signaling, and PKC is a regulator of broader MAPK signaling and not specific to Erk½. Nonetheless, we observed similar effects on the amplification of centrioles (Fig. 3, G to J, and fig. S3, E and G). We think these discrepancies could be due to apical expansion being a cell non-autonomous process, whereas centriole amplification is cell autonomous.

**Figure 3.**
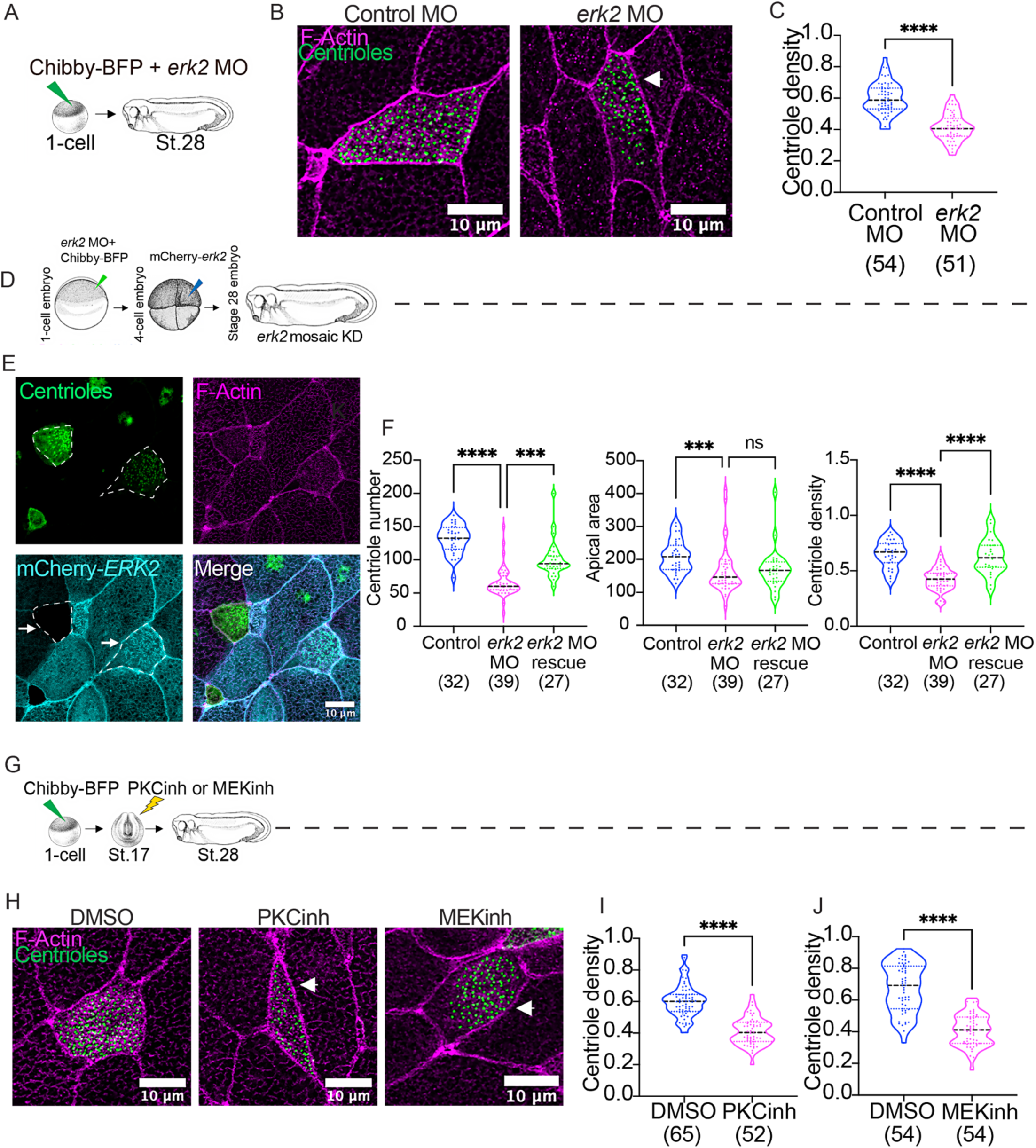
Erk2 is essential for centriole amplification in *Xenopus* MCCs. A. Schematic of the experimental design for either *erk2* knockdown via morpholino (*erk2* MO). Embryos are injected at 1-cell stage with Chibby-BFP (green) and the MO. Embryos are collected at stage 28 for immunostaining and visualization. B. Representative IF images of stage 28 MCCs that are either injected with the standard control (Control MO) or *erk2* morpholino (*erk2* MO), expressing Chibby-BFP (centrioles, green) and are co-stained with Phalloidin (Magenta). Scale bar: 10um. C. Plot depicts comparison of centriole density per MCC between control and *erk2* KD (*erk2* MO) stage 28 embryos. Mann-Whitney non-parametric t-test was used for statistical analysis (****p<0.0001). Data shown is from three independent experiments; number of MCCs from at least 5 embryos per group per experiment is indicated in parentheses. D. Schematic representing the experimental design for mosaic rescue of *erk2*. Embryos were injected with Chibby-BFP and the *erk2* MO at 1-cell stage, followed by injection with mCherry-*erk2* (cyan) in one of the four cells at the 4-cell stage to trace the mosaic rescue, and then allowed to develop until stage 28. E. Representative IF images show mosaic expression of mCherry-*erk2* in MCCs (cyan). The KD effect is indicated by the absence of the cyan signal and the rescue, by the presence of the mCherry-*erk2* localization in the MCC (indicated by arrows in the image sub-panel). Stage 28 embryos were expressing Chibby-BFP (Centrioles, green) and co-stained with Phalloidin (F-Actin, magenta). Scale bar is 10 um. F. Plot depicts a comparison of centriole number, apical area, and centriole density (centriole number/apical area) per MCC for uninjected control (Control), *erk2*-depleted (*erk2* MO), and rescue (*erk2* MO rescue) groups. Kruskal-Wallis test followed by multiple comparisons was performed for statistical analysis between groups (p>0.05 (ns), ***p<0.001, and ****p<0.0001). Data shown is from three independent experiments; the number of MCCs from at least 5 embryos per trial is indicated in parentheses below each group. G. Schematic of the experimental design for either PKC or MEK inhibition. Embryos were injected at 1-cell stage with Chibby-BFP (green) followed by treatment with either PKC inhibitor (25 uM) or MEK inhibitor (25 uM) at stage 17. The untreated controls and treated embryos were collected at stage 28 for immunostaining and visualization. H. Representative IF images of stage 28 MCCs that are either vehicle-treated (DMSO) or treated with PKC inhibitor (PKCinh), or MEK inhibitor (MEKinh), expressing Chibby-BFP (centrioles, green) and are co-stained with Phalloidin (Magenta). Scale bar: 10um. I. The plot depicts a comparison of centriole density per MCC between control and PKC inhibitor-treated stage 28 embryos. Mann-Whitney non-parametric t-test was used for statistical analysis (****p<0.0001). The data shown is from three independent experiments; the number of MCCs from at least 5 embryos per trial is indicated below each group. Plot depicts comparison of centriole density per MCC between control and MEK inhibitor-treated stage 28 embryos. Mann-Whitney non-parametric t-test was used for statistical analysis (****p<0.0001). Data shown is from three independent experiments, the number of MCCs from at least 5 embryos per trial is indicated in parentheses below each group.

### Piezo1 stimulation activates YAP1 in MCCs via the Erk½ pathway

Given the emerging role of Piezo1-mediated calcium influx, Erk and YAP1 activation in promoting cell proliferation and differentiation (*15, 16*), and the importance of YAP1 in the *Xenopus* epidermis during epithelialization and MCC differentiation (*7*), we next sought to dissect the interplay between Piezo1, YAP1, and Erk in regulating centriole number in mature MCCs. As a first step, we examined YAP localization in MCCs using a YAP antibody that detects the active YAP conformation. Compared to vehicle-treated controls where YAP is absent from MCC nuclei, Piezo1 stimulation by Yoda1 activates YAP1 and leads to its nuclear entry in MCCs (fig. S4, A to C). To characterize the dynamic nuclear-cytoplasmic shuttling of YAP1, we used a WT *yap1*-GFP construct previously used by Dr. Lance Davidson (*7*). As expected, Yap1 is localized to the cytoplasm, but to our surprise, we also found that Yap1 localizes to the subapical actin in MCCs (*36*) (Fig. 4, A). Consistent with the results obtained from immunofluorescence staining with the active-YAP1 antibody (fig. S4, A to C), Yoda1- mediated activation of Piezo1 increased nuclear localization of *yap1*-GFP, as shown by a higher nuclear/cytoplasmic ratio (Fig. 4, B and C).

**Figure 4.**
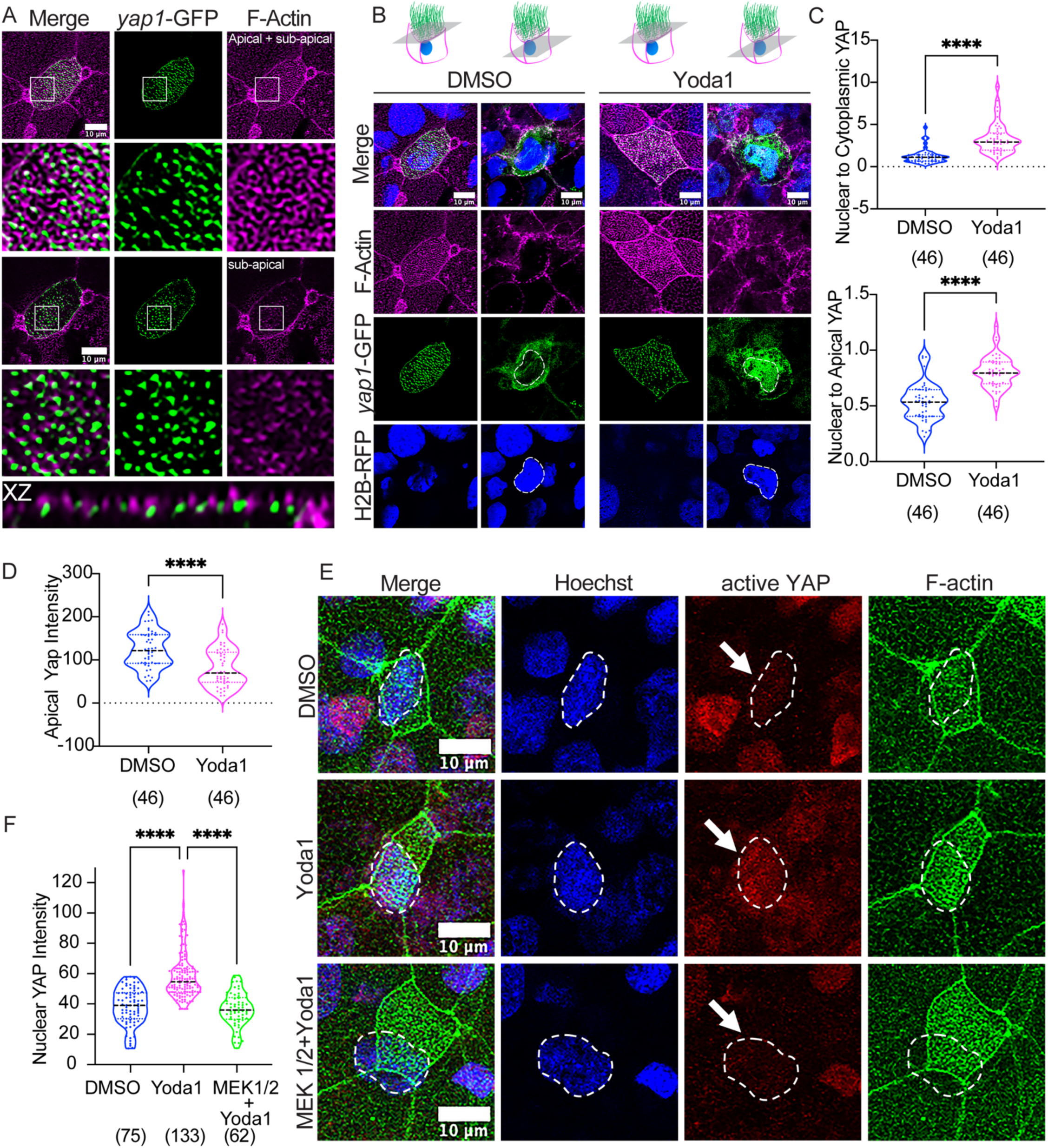
Piezo1 stimulation activates YAP1 in *Xenopus* MCCs via the Erk½. A. Representative images show the localization of Yap to the sub-apical region of the actin network in stage 28 MCCs. Embryos were stained with Phalloidin (magenta) and were injected with *yap1*-GFP (150 pg, green) at 1-cell stage. The XZ cross-section indicates co-localization of YAP with the sub-apical region of actin in MCCs. B. Representative images show the dynamic translocation of Yap in MCCs using a *yap1*-GFP construct, in response to Piezo1 activation via Yoda1. Embryos were injected with *yap1*-GFP (150 pg, green) and a nuclear tracer, *h2b*-RFP (100 pg, blue) at 1-cell stage and treated with either the vehicle control, DMSO, or Yoda1 (25uM) for ten minutes when stage 28 was reached. Apical and nuclear cross-sections were taken through z-stacks and are shown to indicate *yap1*-GFP localization with treatments. XZ cross-sections were also obtained for the control and yoda1 treatment groups to observe the localization of Yap to actin and the nucleus. Embryos were stained with Phalloidin (magenta) to visualize actin. Scale bar is 10 um. C. Plots indicate the ratio of nuclear Yap to cytoplasmic Yap (*Top:* Nuclear to Cytoplasmic YAP) and the ratio of Yap localized to the nucleus versus that localized to apical actin (*Bottom*: Nuclear to Apical YAP) for IF studies in (B). Confocal z-stacks were obtained for each image; for nuclear intensity measurements, the nuclear z-stack sections were extracted, and the image was further processed for intensity measurements in Fiji using the polygon tool. For the apical actin localization measurements, z-stack sections pertaining to the apical and sub-apical regions of the cell were extracted and analyzed for *yap1*- GFP intensity using the polygon tool in Fiji. Additionally, cytoplasmic YAP intensity values were obtained by subtracting *yap1*-GFP nuclear intensity values from both, apical actin Yap and whole cell Yap intensity measurements. Mann-Whitney non-parametric t-test was used for statistical analysis (****p<0.0001). Data shown is from three independent experiments; the number of MCCs from at least 3 embryos per trial is indicated in parentheses below each group. D. Plot indicates the intensity of Yap localized to apical actin in MCCs in stage 28 embryos that were either treated with the vehicle control (DMSO) or Yoda1 (25uM) for ten minutes. Mann-Whitney non-parametric t-test was used for statistical analysis (****p<0.0001). Data shown is from three independent experiments; the number of MCCs from at least 3 embryos per trial is indicated in parentheses below each group. E. Representative IF images show the effect of Erk ½ pathway inhibition via Mek ½ inhibitor (25 uM) on Yap activation in MCCs from stage 28 embryos. Embryos were stained for active-Yap1 (active-YAP, red) and co-stained with Phalloidin (F-Actin, green) and Hoechst (nuclei, blue). Scale bar is 10 um. F. Plot shows the intensity of nuclear Yap1 (active-YAP) per MCC across the different treatment groups in (E). Embryos were pre-treated with Mek ½ inhibitor alone for 1 hour, followed by Mek ½ inhibitor and Yoda1 for 10 minutes (MEK ½ +Yoda1), and compared against embryos treated with Yoda1 alone for 10 minutes (Yoda1, 10 min) and the vehicle control (DMSO). Kruskal-Wallis test followed by multiple comparisons was performed for statistical analysis between groups (p>0.05 (ns), and ****p<0.0001). Data shown is from three independent experiments; the number of MCCs from at least 5 embryos per trial is indicated in parentheses below each group.

Localization of Yap1 to the subapical actin network in multiciliated cells is a novel finding that raises an interesting question - why does Yap1 accumulate at the sub-apical actin network? In a low-tension environment, the Hippo pathway via Lats1/2 phosphorylates Yap1 at serine residues to inhibit its nuclear localization and promote cytoplasmic retention by binding to the 14-3-3 complex or degradation through ubiquitination (*37–41*). One possible explanation is that Yap1 is harvested at the subapical region of the intricate actin meshwork of MCCs for immediate access to mechano-sensation and translocates to the nucleus in response to increased tension. Consistent with this hypothesis, we see that Yoda1-mediated activation of Piezo1 treatment causes an increase in the nuclear/apical ratio of Yap1 and a concurrent significant reduction in the apical enrichment of Yap1 (Fig. 4, B to D).

Whether Yap sequestered by apical actin is essential for tension-dependent centriole amplification is unclear because apical actin enrichment depends on basal body amplification and docking in *Xenopus* and mouse airway cells (Unpublished data; Dr. Mahjoub). There are two possibilities. First, the initial amplification of 75-100 centrioles can generate enough apical F-actin enrichment to sequester Yap, which then can translocate to the nucleus to promote centriole amplification in a tension-dependent manner (*40*). Second, apical actin-sequestered Yap is not essential in the centriole amplification process during MCC morphogenesis. Only cytoplasmic Yap is utilized during tension-dependent centriole amplification. In the future, we plan to address this question by examining how Yap is sequestered by apical actin.

How does Piezo1 activate Yap1 in MCCs? A recent study showed that the Piezo1-Mapk axis activates Yap1 and promotes cell proliferation in hepatocellular carcinoma cells (*17*). Therefore, we hypothesized that MCCs activate Yap1 via the Piezo1-Mapk signaling axis. To test the hypothesis, we activated Piezo1 using Yoda1 in the presence of Mek½ inhibitor or *erk2* MO to block Erk activity. Strikingly, in both cases, Yap1 activation and nuclear entry were significantly diminished (Fig. 4, E and F, and fig. S4, D to F), demonstrating the role of the Piezo1- Mapk axis in regulating Yap1 activity in MCCs. The crosstalk between the Mapk and Hippo cascades in MCCs is intriguing. Studies accounting for cell density regulation in epithelial cells highlight the tension-dependent role of the Ajuba family of proteins in mediating Yap1 activity through the Mapk pathway (*14, 42*). Therefore, it is plausible that MCCs repurpose this crosstalk in response to tension to promote Yap nuclear translocation and regulation of downstream genes essential for centriole amplification.

### Yap1 is essential for centriole amplification in MCCs

We next examined if Yap1 is essential for centriole amplification in MCC by depleting Yap1 using the morpholino oligonucleotide (*yap1* MO). We observed multiple interesting phenotypes suggesting that Yap plays a broader role in MCC morphogenesis. Loss of Yap1 significantly reduced centriole number and density while increasing the apical area (Fig. 5, A to C, and fig. S5, A and B). However, these cells were rounder, suggesting a loss of tension. Consistently, we noticed a significant decrease in apical actin in MCCs, suggesting that Yap1 is essential for actin remodeling in MCCs (Fig. 5, B, and D). We did not measure F-actin in non-MCCs. Next, we verified *yap1* MO specificity using a mosaic rescue experiment. We injected *yap1* MO at the 1-cell stage to deplete Yap from the whole embryo and injected WT *yap1*-GFP in 1 of the 4 cells at the 4-cell stage to rescue centriole amplification and actin in MCCs that receive *yap1*-GFP (Fig. 5E). We noticed that the intensity of the Chibby signal is sometimes higher in Yap KD cells. While we do not completely understand the underlying reason, one possibility is that cells with fewer centrioles have excess Chibby protein available that accumulates at the centrioles. Nonetheless, we show that *yap1*-GFP rescued centriole density, centriole number, and apical actin in MCCs, confirming the role of Yap1 in centriole number control and actin remodeling in MCCs (Fig. 5, F to H, and fig. S5, C and D).

**Figure 5.**
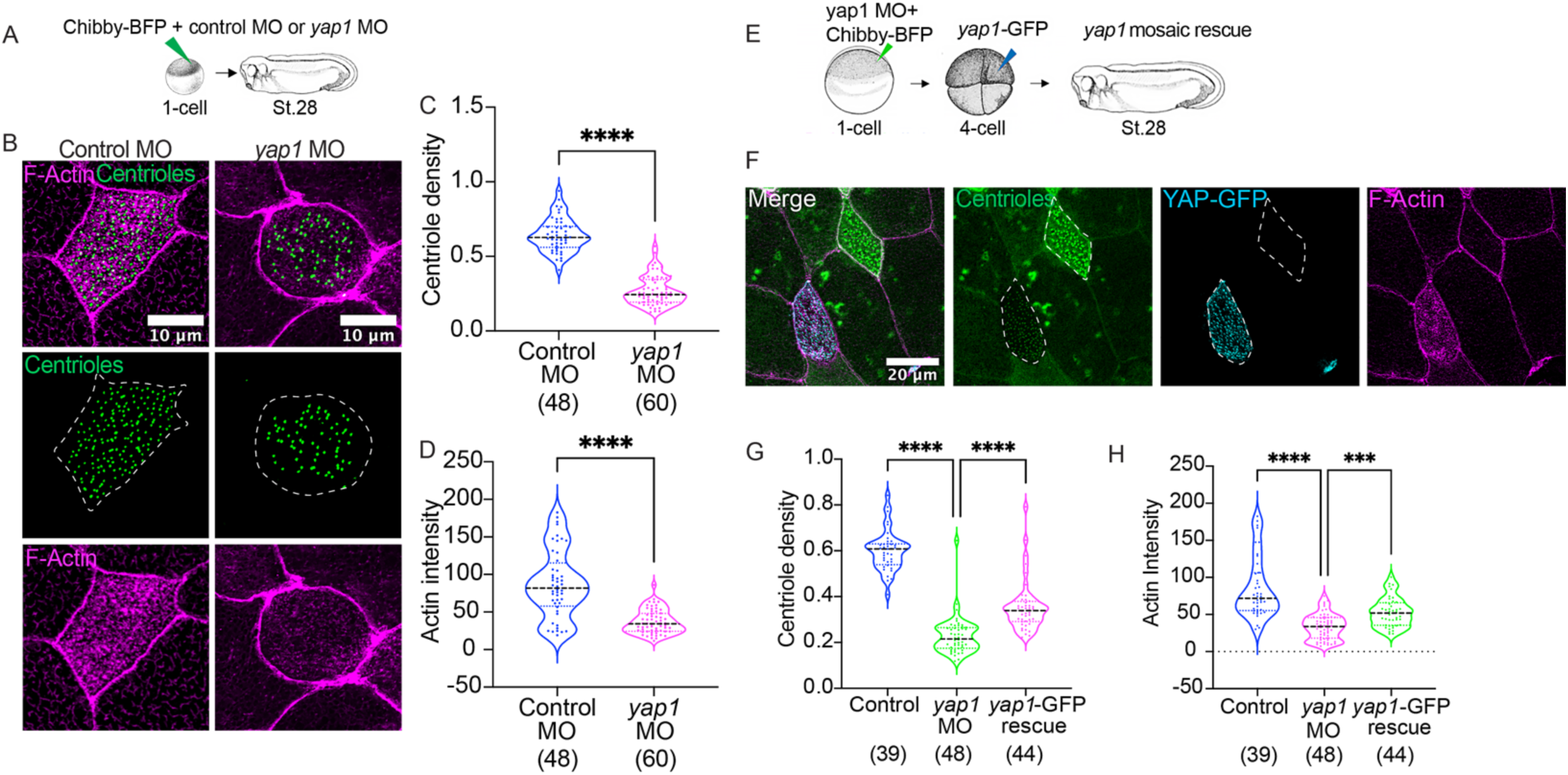
Yap1 is essential for centriole amplification and apical actin enrichment in *Xenopus* MCCs. A. Schematic of the experimental design for assessing centriole number in *yap1* knockdown MCCs via morpholino (*yap1* MO). Embryos are injected at 1-cell stage with Chibby-BFP (green) and either the standard control morpholino (Control MO) or yap1 morpholino (*yap1* MO). Embryos are collected at stage 28 for immunostaining and visualization. B. Representative IF images of stage 28 MCCs that are either injected with the standard control (Control MO) or *yap1* morpholino (*yap1* MO), expressing Chibby-BFP (centrioles, green) and are co-stained with Phalloidin (magenta). Scale bar: 10um. C. Plot depicts comparison of centriole density per MCC between standard control (Control MO) and *yap1* KD (*yap1* MO) stage 28 embryos. Mann-Whitney non-parametric t-test was used for statistical analysis (****p<0.0001). Data shown is from three independent experiments; the number of MCCs from at least 5 embryos per trial is indicated in parentheses below each group. D. Plot depicts comparison of actin intensity per MCC between standard control (Control MO) and *yap1* KD (*yap1* MO) stage 28 embryos. Mann-Whitney non-parametric t-test was used for statistical analysis (****p<0.0001). Data shown is from three independent experiments; the number of MCCs from at least 5 embryos per trial is indicated in parentheses below each group. For actin intensity measurements, the Fiji polygon tool was used to extract the medial actin region, and intensity measurements were made in Fiji. Data shown is from three independent experiments; the number of MCCs from at least 5 embryos per trial is indicated in parentheses below each group. E. Schematic of the experimental design for assessing mosaic rescue of *yap1* knockdown MCCs (via *yap1* MO) using the WT *yap1*-GFP construct. Embryos were injected at 1-cell stage with Chibby-BFP (green) and *yap1* MO. Embryos were then injected with *yap1*-GFP in one of the four cells at the 1 in 4 cell stage to obtain mosaic rescue and were collected at stage 28 for immunostaining and visualization. F. Representative IF images of stage 28 MCCs that are either depleted of *yap1* via MO (*yap1* MO) or rescued by the WT *yap1*-GFP construct (via *yap1*-GFP, cyan), expressing Chibby-BFP (centrioles, green) and co-stained with Phalloidin (magenta). Scale bar: 10um. Plot depicts comparison of centriole density per MCC between unperturbed controls, yap1 KD (*yap1* MO), and *yap1*-rescued stage 28 MCCs (yap1-GFP rescue). Kruskal-Wallis test followed by multiple comparisons was performed for statistical analysis between groups (***p<0.001, and ****p<0.0001). Data shown is from three independent experiments; the number of MCCs from at least 5 embryos per trial is indicated in parentheses below each group.

Plot depicts comparison of actin intensity per MCC between unperturbed controls, yap1 KD (*yap1* MO), and *yap1*-rescued stage 28 MCCs (yap1-GFP rescue). Kruskal-Wallis test followed by multiple comparisons was performed for statistical analysis between groups (***p<0.001, and ****p<0.0001). Data shown is from three independent experiments; the number of MCCs from at least 5 embryos per trial is indicated in parentheses below each group.

### Yap1 and Piezo1 regulate Foxj1 expression in MCCs

How does Yap regulate centriole amplification? One possibility is that Yap may regulate centriole amplification via its role in regulating the apical actin enrichment in MCCs. However, if and how F-actin enrichment links to centriole amplification is not known. Another possibility is that Yap may act as a co-transcriptional activator to control the transcription of genes essential for centriole amplification. Interestingly, a study in adult murine trachea indicated that Yap1 is necessary for the expression of Foxj1, a master regulator of motile ciliogenesis, and thus, the assembly of multiple cilia in differentiating MCCs (*43*). However, this study did not examine its effect on centriole amplification. Moreover, Yap is also shown to be essential for motile ciliogenesis in the left-right organizer via regulating Foxj1 expression (*44*), suggesting that Yap may regulate Foxj1 expression in ciliated cells.

While the relation between Yap and Foxj1 has been shown before, it is possible that Yap may also regulate other key genes in centriole amplification. Therefore, we decided to examine the expression of four genes, *foxj1*, *myb*, *plk4*, and *ccno*, essential for centriole amplification and multiciliogenesis using the HCR-RNA FISH in Yap morphants (*12, 45–48*). We examined stage 25 because, from stages 22-23 onwards, the embryo undergoes a significant change in shape from round to elongated, exerting more tension on the multiciliated epithelium. Related to this process, most MCCs finish apical insertion and begin to undergo apical expansion, which causes tension-dependent centriole amplification. Unfortunately, we could not detect a meaningful RNA signal in the genes *plk4* and *ccno,* but we did observe a strong signal in *myb* and *foxj1* in the MCCs of control embryos (fig. S6, A and B). Both *myb* and *foxj1* signals were significantly reduced in Yap morphants suggesting the role of Yap in regulating transcriptional regulation of centriole amplification.

Given the previous reports on the Yap regulation of Foxj1 and our HCR-RNA FISH results, we next confirmed a reduction in *foxj1* using whole mount *in situ* hybridization assay. We injected *yap1* MO in one of the 2-cell stage embryos. We compared *foxj1* RNA expression between uninjected control and MO-injected sides of the same embryo. We found that depletion of Yap1 resulted in a significant decrease in *foxj1* mRNA expression (Fig. 6A). Because Piezo1 regulates Yap1 activation, we also examined whether Piezo1 regulates *foxj1* expression. Like Yap1 depletion, loss of Piezo1 using *piezo1* MO also significantly decreased *foxj1* mRNA levels (Fig. 6B), suggesting an essential role for mechanosensitive proteins in regulating Foxj1 levels in *Xenopus* MCCs.

**Figure 6.**
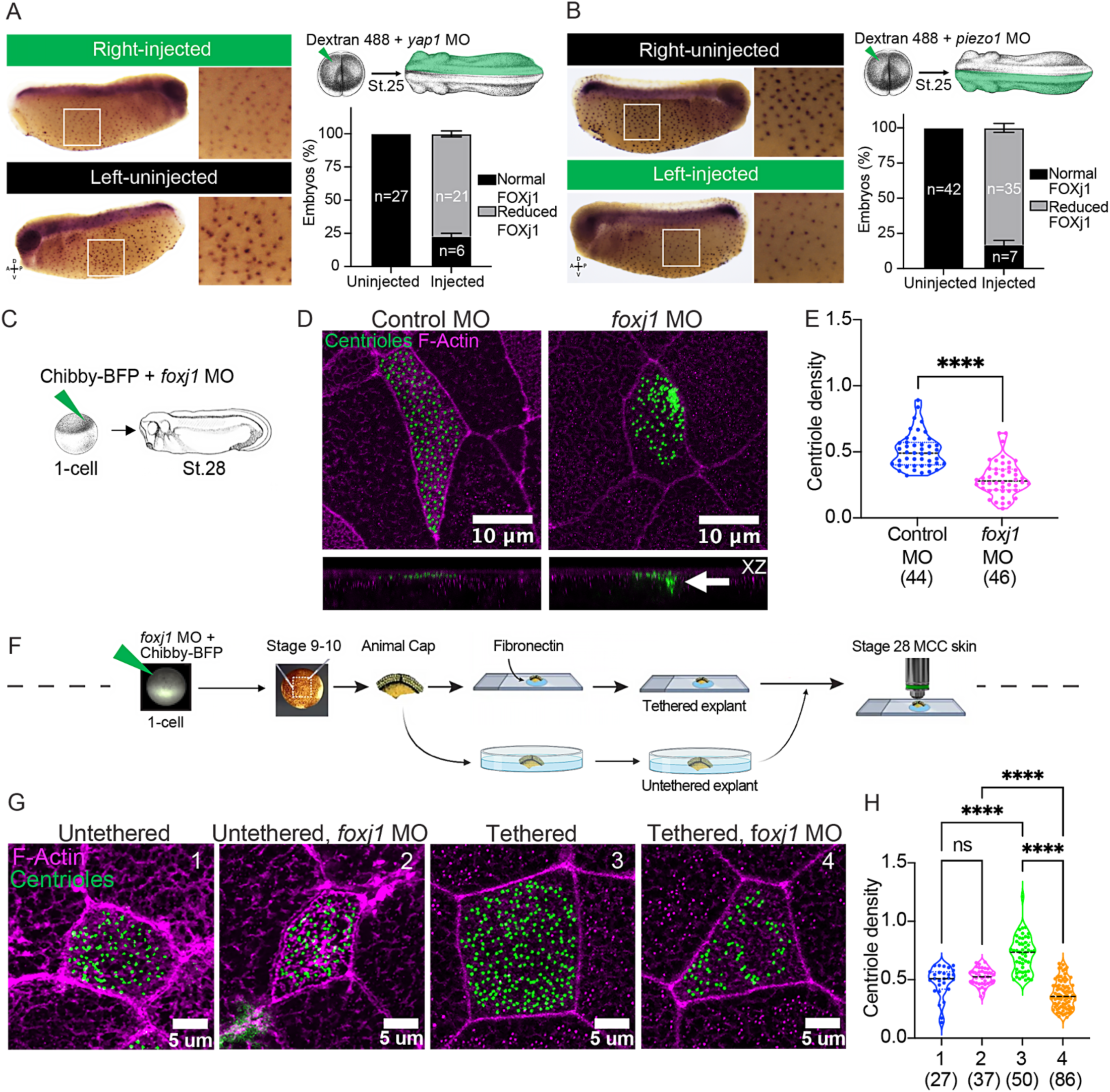
Foxj1, downstream of Yap1, regulates centriole amplification in a tension-dependent manner. A. Representative images of stage 25 embryos injected with *yap1* MO and a tracer (Dextran-488+*yap1* MO) in one blastomere at the two-cell stage, sorted at stage 25, and probed for foxj1 mRNA expression via whole mount in situ hybridization. *Left*: the top panel indicates the uninjected side of the embryo (black) that serves as control, and the bottom panel shows the injected side (green) which expresses Dextran- 488 and is depleted of Yap1. The bar plot shows the percentage of embryos with normal (black) and reduced (grey) foxj1 mRNA expression (error bars: mean +/- SEM). Data shown is from three independent experiments, and n=number of embryos. B. Representative images of stage 25 embryos injected with *piezo1* MO and a tracer (Dextran-488+*piezo1* MO) in one blastomere at the two-cell stage, sorted at stage 25, and probed for foxj1 mRNA expression via whole mount in situ hybridization. *Left*: the top panel indicates the uninjected side of the embryo (black) that serves as control, and the bottom panel shows the injected side (green) which expresses Dextran-488 and is depleted of Piezo1. The bar plot shows the percentage of embryos with normal (black) and reduced (grey) foxj1 mRNA expression (error bars: mean +/- SEM). Data shown is from three independent experiments, and n=number of embryos. C. Schematic of the experimental design for assessing centriole number in *foxj1* knockdown MCCs via morpholino (*foxj1* MO). Embryos are injected at 1-cell stage with Chibby-BFP (green) and either the standard control morpholino (Control MO) or foxj1 morpholino (*foxj1* MO). Embryos are collected at stage 28. D. Representative IF images of stage 28 MCCs that are either injected with the standard control (Control MO) or *foxj1* morpholino (*foxj1* MO), expressing Chibby-BFP (centrioles, green) and are co-stained with Phalloidin (magenta). Scale bar: 10um. E. Plot depicts comparison of centriole density per MCC between control and foxj1 KD (*foxj1* MO) stage 28 embryos. Mann-Whitney non-parametric t-test was used for statistical analysis (****p<0.0001). Data shown is from three independent experiments; the number of MCCs from at least 5 embryos per trial is indicated in parentheses below each group. F. Schematic representing the experimental design for animal explant extraction of either control or *foxj1* KD embryos prior to staining and visualization. Animal explants were surgically extracted at stage 9-10 from embryos that were injected with Chibby-BFP (green) and *foxj1* MO at 1-cell stage, followed by culture in either fibronectin-coated slides (tethered) or an uncoated dish with media (untethered). The explants were then allowed to develop until stage 28. The explants were stage-matched with unmanipulated sibling embryos. G. Representative IF images of stage 28 MCCs in animal explants expressing Chibby-BFP (centrioles, green) and co-stained with Phalloidin (Magenta). The tethered explant depleted of foxj1 (4: Tethered, *foxj1* MO) is compared against the uninjected tethered control (3: Tethered) and the untethered groups (Untethered (1) and Untethered, *foxj1* MO (2)). Scale bar is 10um. Plot shows comparison of the centriole density per MCC between the different groups in (G) that indicates a reduction in centriole density with foxj1 depletion even in the presence of mechanical tension. Kruskal-Wallis test followed by multiple comparisons was performed for statistical analysis between groups (p>0.05 (ns), ***p<0.001, and ****p<0.001). Data shown is from three independent experiments; the number of MCCs from at least 5 embryos per trial is indicated in parentheses below each group.

### Foxj1 regulates centriole number in a tension-dependent manner

Myb was previously reported to be essential for centriole (*47*), but Foxj1 was a new candidate, which piqued our interest, so we decided to test the role of Foxj1 in centriole number control. Foxj1 depletion in *Xenopus* showed impaired ciliogenesis, specifically a reduced number of motile cilia in the left-right organizer and the multiciliated epidermis (*46*). The Kintner group showed that a reduced number of motile cilia is associated with fewer centrioles in *Xenopus* MCCs. However, they did not quantify the number of centrioles. Therefore, we decided to quantify the number of centrioles in MCCs of Foxj1-depleted MCCs.

We depleted Foxj1 using a translation-blocking morpholino at the 1-cell stage and counted the number of centrioles in MCCs using IMARIS 3D reconstruction of MCCs, as shown previously in our Elife paper (*11*). Albeit some centrioles appeared to remain undocked, consistent with prior reports (*49*), we found that morpholino-based knockdown of Foxj1 blocked the complete amplification of centrioles in comparison to the control embryos (Fig. 6, C to E, and fig. S6, C and D). While Foxj1 depletion resulted in significantly fewer centrioles than the controls, the apical area of MCCs remained unaffected, reducing centriole density. These results are interesting when compared to Yap1 KD. Foxj1 loss and Yap1 loss both lead to reduced centrioles and F-actin enrichment in MCCs, but only Yap KD leads to changes in the MCC size and shape. We hypothesize that Foxj1 is expressed only in MCCs, whereas Yap is expressed in MCCs and non-MCCs. Thus, the loss of Yap has broader consequences, leading to a loss of global tension in the epithelium, resulting in changes in MCC size and shape.

Our results appear to contradict the previous studies in mice with a Foxj1 knockout (*50*). Specifically, Foxj1 knockout mice show impaired ciliogenesis and a reduced number of motile cilia. Transmission electron microscopy analysis clearly shows centrioles that fail to dock at the apical surface, and this defect in centriole docking is attributed to impaired ciliogenesis and a reduced number of motile cilia. While we agree that the defect in centriole docking as a cause for impaired ciliogenesis is evident, these studies did not count the number of centrioles directly. Thus, these studies have not addressed whether Foxj1 is also essential for centriole amplification, leading to a reduced number of motile cilia. Second, we presume that the gene functions in the MCC morphogenesis process are the same or similar between the mouse airway and the *Xenopus* epidermis. However, there is a clear distinction between the MCC morphogenesis process in these two model systems. Mouse airway MCCs are specified *de novo* in the superficial epithelium and undergo moderate apical expansion. *Xenopus* epidermal MCCs, on the contrary, undergo radial intercalation from the basal cell layer to the superficial epithelium, followed by expansion of their apical surface from 0 to 300^2^ μm. These dramatic differences in MCC morphogenesis may also explain differences in the adaption of gene function in mice and frogs.

*Foxj*1 expression is regulated by a mechanotransduction molecule Yap (*43*) and is required for motile cilia formation in response to strain in *Xenopus* left-right organizer (LRO), indicating that Foxj1 could be a mechanosensitive transcriptional regulator of motile ciliogenesis (*51*). Therefore, we investigated whether Foxj1 regulates centriole amplification in a tension-dependent manner. We used our established system of tethered (under tension) and untethered *Xenopus* stem cell explants to test the role of Foxj1 in centriole amplification. Depletion of Foxj1 using MO did not affect centriole density, centriole number, and apical area in untethered caps (Fig. 6, F to H, and fig. S6, E and F). However, centriole number and centriole density both were significantly affected in tethered explants, demonstrating a novel mechanosensitive role for Foxj1 in regulating centriole number and centriole density in MCCs (Fig. 6, F to H, and fig. S6, E and F). How does Foxj1 regulate centriole amplification in a tension-dependent manner? Actin cytoskeleton plays a major role in mechanotransduction in epithelial cells under tension, and Foxj1 is essential for F-actin remodeling and enrichment in MCCs (*52*), suggesting a hypothesis that Foxj1-regulated actin remodeling is essential for tension-dependent centriole amplification in MCCs. The second possibility is that Foxj1 is a transcription factor and regulates the expression of genes essential for centriole amplification in a tension-dependent manner.

### Piezo1 correlates with centriole amplification during MCC development

Piezo1 localizes at the cell junctions and is mobile at the apical surface of many epithelial cells (*53*). However, in MCCs, Piezo1 co-localizes near centrioles (fig. S7A). Using super-resolution microscopy, we show that it forms a ring-shaped structure with an outer diameter of approximately 250nm (Fig. 7A). When we counted the number of Piezo1 puncta (likely consisting of many Piezo1 molecules) and the number of centrioles during the early stages of MCC development, from stages 18 to 28, we saw a robust correlation, suggesting that the transcriptional program that regulates centriole amplification may also regulate Piezo1 expression (fig. S7B). Consistent with our hypothesis, HCR RNA-FISH analysis showed increased Piezo1 mRNA levels during MCC differentiation (from stage 18 to 25) and a decrease around stage 28 when MCCs mature (fig. S7, C, and D).

**Figure 7.**
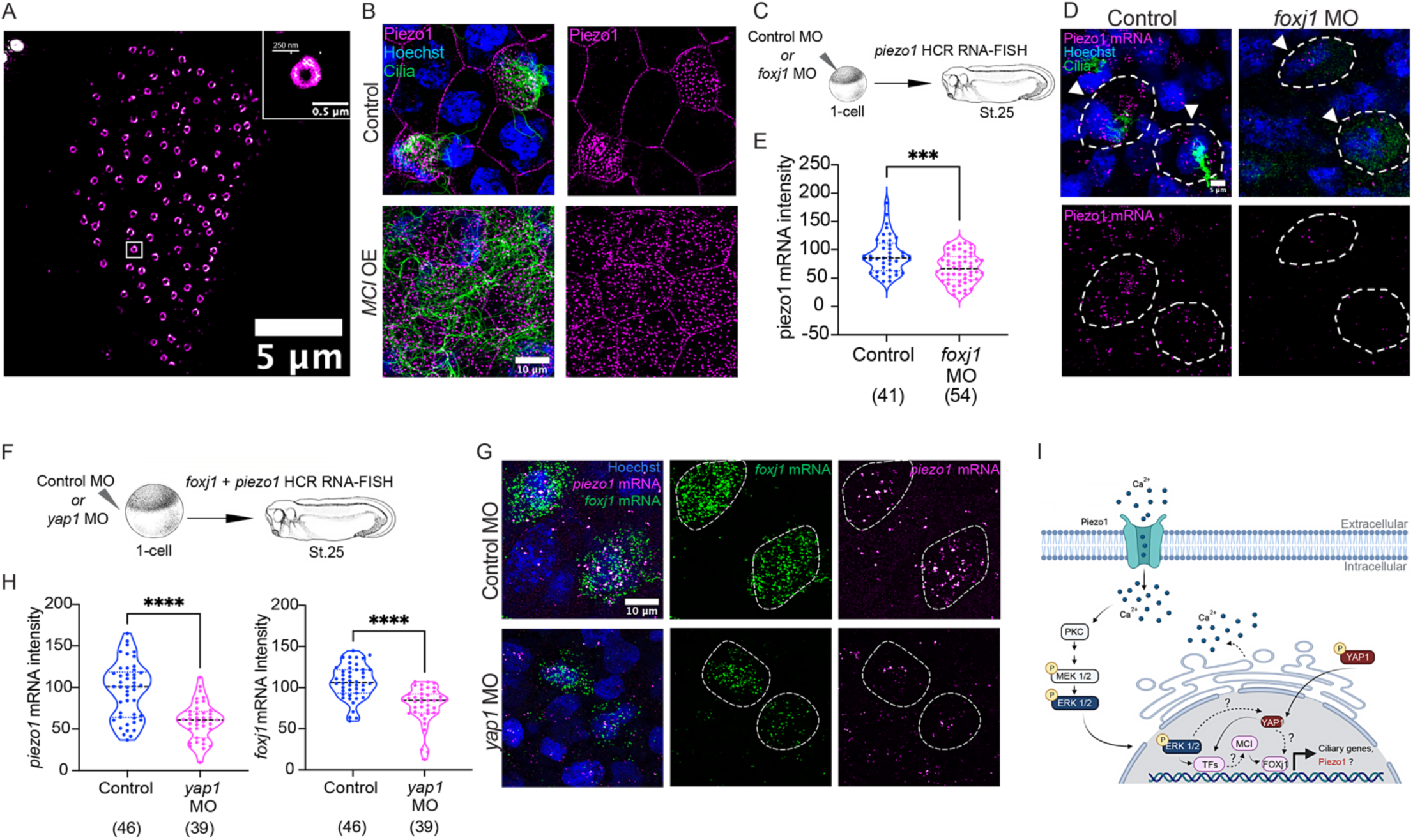
YAP1-Foxj1 signaling regulates Piezo1 expression in *Xenopus* MCCs in a positive feedback loop. A. A super-resolution image of a mature multiciliated cell (MCC) from a stage 28 embryo was immuno- stained with Piezo1 antibody (magenta). Scale bars: 5 um and 1.5 um (inset). B. Schematic of the experimental design for assessing piezo1 puncta expression with over-expression of multicilin (*MCI OE*). Embryos are injected at the 1-cell stage with multicilin (*MCI*, 50 pg) and induced for multicilin expression via dexamethasone treatment (0.1 mg/ml) at stage 10. Untreated embryos serve as controls and the treated embryos are collected at stage 28. Representative IF images of stage 28 MCCs that are either uninduced (Control) or induced for Multicilin over-expression (*MCI* OE) are stained for Piezo1 (magenta), and cilia (acetylated tubulin, green). Nuclei are counterstained with Hoechst (blue). Scale bar: 10um. C. Schematic of the experimental design for *piezo1* mRNA expression in foxj1 knockdown MCCs via morpholino (*foxj1* MO). Embryos are injected at 1-cell stage with either the standard control morpholino (Control MO) or *foxj1* morpholino (*foxj1* MO). Embryos are collected at stage 25 for HCR RNA-FISH and visualization. D. Representative HCR RNA-FISH images of stage 25 MCCs that are either injected with the standard control (Control MO) or *foxj1* morpholino (*foxj1* MO). Embryos are probed with Piezo1 (*piezo1* mRNA, magenta) HCR RNA probes and stained for cilia (acetylated tubulin, green). Nuclei are counterstained with Hoechst (blue). Scale bar: 10um. E. The plot shows a comparison between control and Foxj1 KD (*foxj1* MO) stage 25 MCCs for *piezo1* mRNA intensity per MCC. Mann-Whitney non-parametric t-test was used for statistical analysis (****p<0.0001). Data shown is from three independent experiments; the number of MCCs from at least 5 embryos per group per trial is indicated in the parentheses below each group. F. Schematic of the experimental design for Piezo1 and Foxj1 mRNA expression in *yap1* knockdown MCCs via morpholino (*yap1* MO). Embryos are injected at 1-cell stage with either the standard control morpholino (Control MO) or yap1 morpholino (*yap1* MO). Embryos are collected at stage 25 for HCR RNA-FISH and visualization. G. Representative HCR RNA-FISH images of stage 25 MCCs that are either injected with the standard control (Control MO) or *yap1* morpholino (*yap1* MO). Embryos are probed with piezo1 (*piezo1* mRNA, magenta) and foxj1 (*foxj1* mRNA, green) HCR RNA probe. Nuclei are counterstained with Hoechst (blue). Scale bar: 10um. Plots show a comparison between control and Yap1 KD (*yap1* MO) stage 25 MCCs for *piezo1* mRNA intensity (*left*), and *foxj1* mRNA intensity (*right*) per MCC. Mann-Whitney non-parametric t-test was used for statistical analysis (****p<0.0001). Data shown is from three independent experiments; the number of MCCs from at least 3 embryos per group per trial is indicated in the parentheses below each group.

### YAP1-Foxj1 signaling regulates Piezo1 expression in a positive feedback loop

We performed a series of experiments to test whether a transcriptional program that regulates centriole amplification also regulates Piezo1 expression. Since Piezo1 colocalizes with centrioles at the apical surface in MCCs, we first examined whether the over-expression of Multicilin, a master regulator of multiciliate cell differentiation and centriole amplification, affected Piezo1 expression in the *Xenopus* epidermis (*49*). To over- express Multicilin, embryos were injected with a glucocorticoid-inducible form of Multicilin, MCI-HGR RNA (*49*), at the 1-cell stage and induced for over-expression with Dexamethasone at the beginning of gastrulation. As expected, over-expression of Multicilin resulted in non-MCCs becoming MCCs, depicted by an increased number of centrioles in each cell (fig. S7, E, MCI OE). Notably, elevated levels of multicilin also increased the number of Piezo1 puncta on the apical surface in all cells (Fig. 7B), suggesting that Multicilin acts not only upstream of centriole amplification but also Piezo1 to regulate its expression. However, when Foxj1 was spatially depleted in the multicilin over-expressed cells via a mosaic knockdown approach, centriole amplification was blocked (fig. S7, E and F), thereby demonstrating that Foxj1 plays a crucial role downstream of MCI in determining the final number of centrioles in MCCs. Since KD of Foxj1 blocks centriole amplification, we next wanted to test whether depletion of Foxj1 affected Piezo1 expression in MCCs. Piezo1 mRNA levels dropped significantly with Foxj1 depletion suggesting a positive feedback loop critical for regulating Piezo1 expression in MCCs (Fig. 7, C to E).

Since Yap1 is critical for regulating Foxj1 expression and centriole amplification in MCCs, and centriole amplification correlates with Piezo1 expression, we hypothesized that depletion of Yap1 will also affect the expression of Piezo1 downstream. To test this idea, we injected embryos with the *yap1* MO at the 1-cell stage and probed for both *foxj1* and *piezo1* mRNA expression levels using HCR-RNA FISH. Notably, Yap1 depletion showed reduced mRNA expression of *piezo1*, suggesting a critical feedback mechanism required for centriole amplification in MCCs (Fig. 7, F to H).

Illustration depicts the proposed Piezo1-mediated PKC-Erk-Yap1 cascade initiated by calcium entry into the cell through the opening of the mechanosensitive ion channel. The cascade in turn activates the master regulators of MCC differentiation and ciliogenesis (multicilin (MCI), and foxj1), including the upregulation of Piezo1 through a positive feedback loop.

## DISCUSSION

Our previous study showed that *Xenopus* epidermal MCCs amplify centrioles in two steps (*11*). In the first step, they synthesize a default number of 75-100 centrioles. In the second step, they amplify centrioles as the apical area increases in response to tension via Piezo1. In this study, we show that Piezo1 activation triggers calcium influx. Calcium, a secondary messenger, activates protein kinase C (PKC), and initiates the Erk ½ / MAPK cascade, activating Yap1 downstream. Yap1 regulates ciliary gene expression and modulates Piezo1 expression via Foxj1 through a positive feedback loop (Fig. 7I). Therefore, the feedback mechanism regulating Piezo1 is critical to understanding this stepwise centriole amplification process. Previous studies have shown that the apical F-actin network absorbs mechanical stress and reduces Piezo1’s sensitivity to mechanical tension (*54*). Thus, with every amplification cycle, MCCs may need to make more Piezo1 to maintain the sensitivity to mechanical tension and amplify centrioles further. However, an alternative possibility is that Piezo1 puncta at ciliary bases are not essential for centriole amplification but to detect local tension generated by the bending of beating cilia. Piezo1 at cell junctions is necessary and sufficient for sensing the tension and inducing centriole amplification.

More research is needed to identify other parallel pathways that may interact with the mechanotransduction cascade in controlling centriole number, apical area, or both. For example, a recent study has implicated the role of Plk4, a master regulator of centriole biogenesis, in driving the centriole amplification correlated to the apical area of MCCs in mouse tracheal epithelial cells (mTECs) (*12*). It is plausible that Plk4 interacts with the identified mechanochemical axis and is essential in the tension-dependent amplification of centrioles in MCCs. Further, other mechanosensitive molecules have been found to localize with apical F-actin and near centrioles in MCCs. For example, focal adhesion proteins FAK, Vinculin, and Paxillin have been shown to regulate ciliogenesis via subapical actin enrichment and basal body organization (*55*). These proteins play a crucial role in mechanotransduction, and both Piezo1 and Yap have been shown to interact with focal adhesion proteins in cycling cells. Whether these proteins play a similar role in MCCs in controlling centriole amplification needs to be explored.

In conclusion, our study ambitiously tackles a critical question in cell biology: How do cells regulate organelle numbers? Here, we demonstrate that post-mitotic MCCs utilize the mechanosensitive cell cycle pathway Piezo1- Erk1/2-Yap1 to control the number of centrioles (Fig. 7I), offering critical insights into pathologies like primary ciliary dyskinesia, reduced generation of multiple motile cilia (RGMC), and cancer, where mutations in Piezo1, Yap1, and Foxj1 have been implicated in aberrant centriole/cilia numbers (*1, 11, 36, 43*). Moreover, we demonstrate Foxj1 as a key mechanosensitive transcription factor and effector of Yap, playing an essential role in tension-dependent centriole amplification. Lastly, MCCs play central roles in the respiratory tract and brain ventricles to circulate the CSF and ovaries to push the ova. Our exploration of organelle number regulation in MCCs also provides critical insights into the etiology of conditions like hydrocephalus (enlargement of brain ventricles due to CSF accumulation) and infertility, highlighting the broader significance of our research.

## MATERIALS AND METHODS

### Key Resource Table

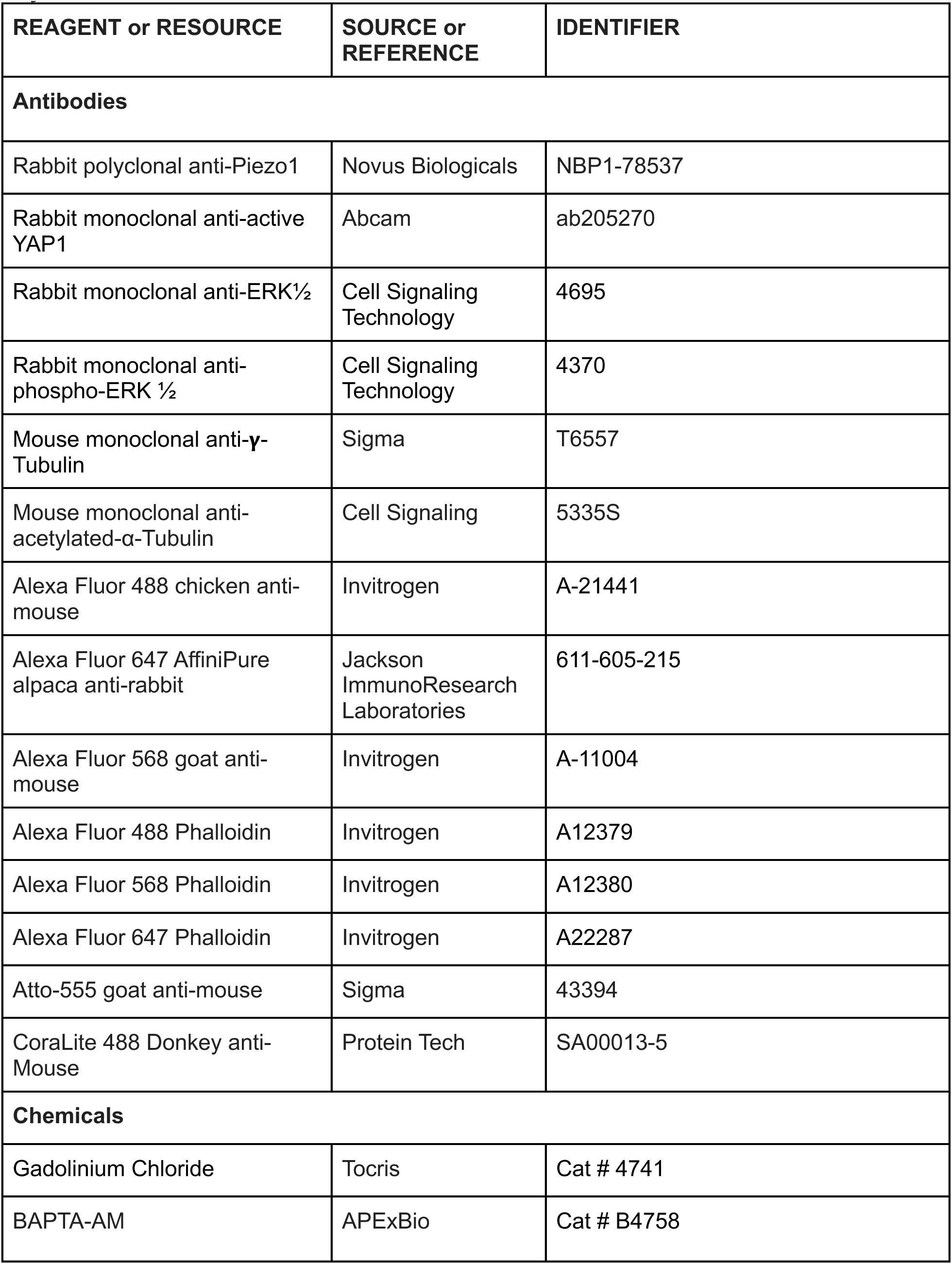
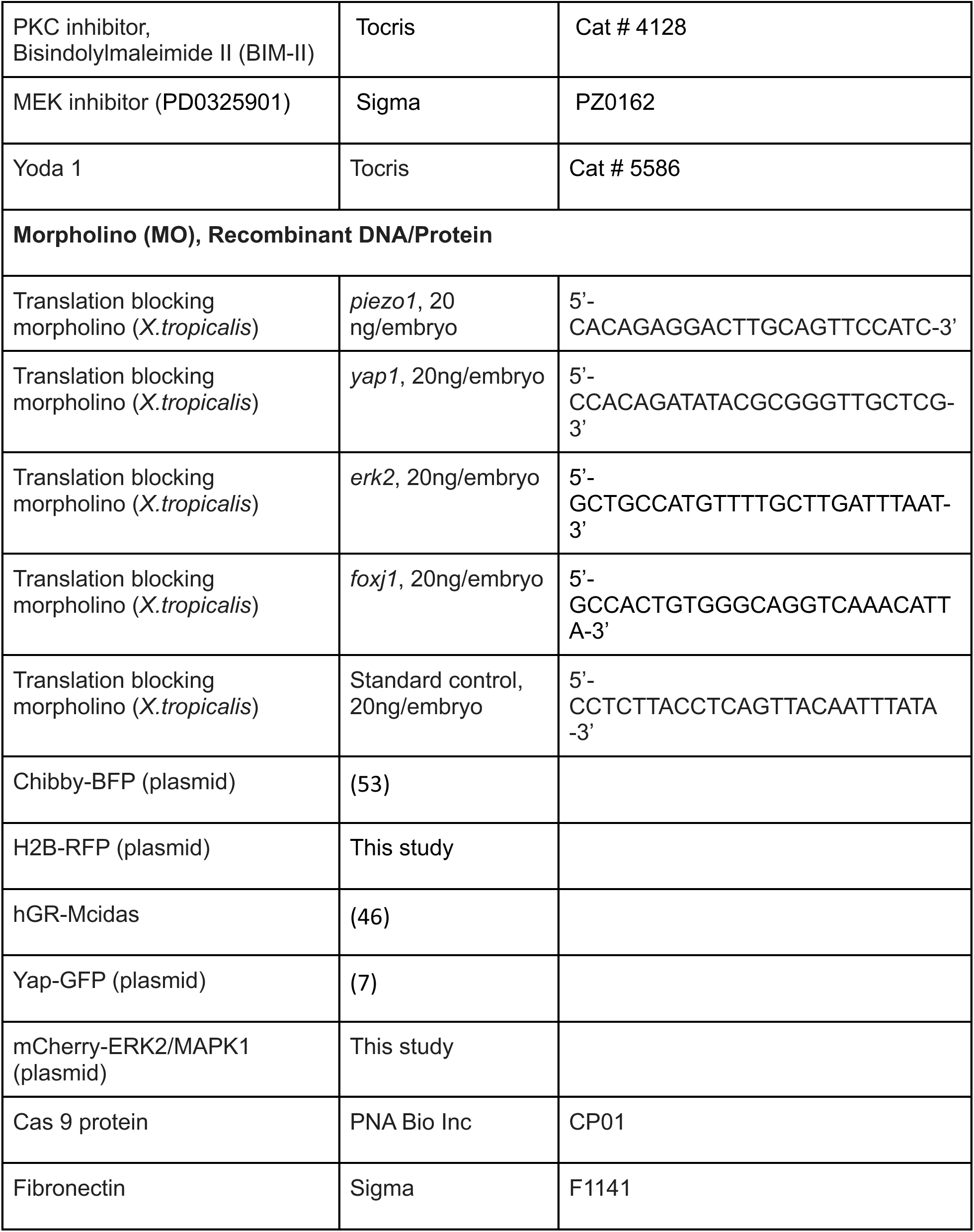

### Animal Husbandry and *In Vitro* Fertilization

*Xenopus tropicalis* were housed and cared for in the animal care facility according to established protocols that were approved by the University of Virginia Institutional Animal Care and Use Committee (IACUC). Embryos were generated via *in vitro* fertilization as previously described (*11, 56*). Briefly, the testes were harvested from an adult male in 1x MBS with 0.2%BSA, crushed, and then incubated with the eggs from an adult female for 3 minutes at room temperature (RT). The fertilized eggs were then flooded with 0.1x MBS (pH 7.8-8) for 10 minutes, followed by the removal of the jelly coat using 3% Cysteine in 1/9x Modified Ringer’s (MR) solution (pH 7.8-8) for 6 minutes. The embryos were then washed with 0.1x MBS (pH 7.8-8), immersed in 2.5% Ficoll, and used for microinjections or allowed to develop in 1/9x MR + 1% gentamycin to the appropriate stages according to the established protocols and previously annotated staging table (*56–59*).

### Animal Explants Dissection

Embryos were collected at the onset of early gastrulation, stage 10, and used for the dissection of animal explants. A fine hair tool was used to cut out a section from the animal pole in a pool of Danilchik’s for Amy (DFA) medium. Following dissection, the tissue was trimmed on the sides to remove the equatorial cells and laid on a glass slide coated with Fibronectin (Sigma, 1mg/ml) immersed in DFA solution. The tissue was then allowed to tether and spread on the fibronectin-coated slides overnight until they reached stage 28. Control embryos were used to stage-match the extracted animal explants.

### Microinjection of mRNAs and Morpholino (MO) Oligonucleotides

*Xenopus tropicalis* embryos were generated by *in vitro* fertilization and raised to appropriate stages in media containing 1/9xMR and gentamycin. Morpholino oligonucleotides and mRNA were injected into one-cell, two-cell, or four-cell embryos with the help of a Picoliter microinjector system (Warner Instruments), as described previously [43]. The following constructs were injected at either 1-cell, 2-cell, or 4-cell stage: piezo1 translation blocking MO (10ng/4-cell embryo, 5’- CACAGAGGACTTGCAGTTCCATC-3’), erk2 translation blocking MO (20ng/1-cell and 10ng/4-cell embryo, 5’- GCTGCCATGTTTTGCTTGATTTAAT-3’), yap1 translation blocking MO (20ng/1-cell, 20ng/2cell, and 10ng/4-cell embryo, 5’-CCACAGATATACGCGGGTTGCTCG-3’), foxj1translation blocking MO (20ng/1-cell embryo, 5’-GCCACTGTGGGCAGGTCAAACATTA-3’), and standard control MO (20ng or 10ng per embryo, 5’-CCTCTTACCTCAGTTACAATTTATA-3’). mRNA for chibby blue fluorescent protein (Chibby-BFP, 100pg), histone-2b red fluorescent protein (H2B-RFP, 100pg), and Dextran-488 (0.8 mg/ul) were injected as markers and tracers as required. In vitro, capped mRNAs were generated with the help of the mMessage machine kit (Ambion). For *yap1* and *erk2* KD-rescue experiments, embryos were first injected with chibby-BFP at the 1-cell stage with the corresponding MO and then with either yap1-GFP (150pg) or mCherry- MAPK1 (150pg) in 1cell at the 4-cell stage. The embryos were then allowed to mature to stage 28 prior to fixation and mounting. The mCherry-MAPK1 construct was generated using Gateway cloning as per the manufacturer’s protocol.

### Generation of mCherry-MAPK1 construct

The mCherry-MAPK1 construct was generated using Gateway cloning as per the manufacturer’s protocol. Briefly, the entry clone (pDONR223-MAPK1-WT, plasmid#82145) was purchased from Addgene and transferred into a destination vector (362 pCS Cherry DEST, plasmid#13075, Addgene) using the LR Clonase II enzyme mix, which is provided with the LR reaction kit. Following incubation at 25C for 1 hour, the reaction wawas terminated by incubation with Proteinase K solution at 37C for 10 minutes. The LR reaction was then transformed into TOP10 competent cells with the appropriate selection marker suited for the destination vector. Following colony selection, the purified plasmid was sequenced and verified prior to RNA synthesis using the mMessage mMachine^TM^ SP6 Transcription Kit.

### Calcium Chelation Assay and Chemical Treatments

BAPTA-AM (APExBio) was prepared at 10mM stock solution. For intracellular calcium chelation experiments, embryos were incubated at embryonic stage 17 with 100uM of BAPTA-AM in 11.1% MR (1/9xMR), with replenishment every two hours. Experiments were also performed with 33% divalent-free (⅓ DVF) medium that lacked calcium chloride and magnesium chloride compared to the MR medium (*60*) to chelate extra-cellular calcium. Similar to BAPTA-AM treatment, embryos or animal caps were incubated at stage 17 with either 30uM of Gadolinium chloride, 25 uM of BIM-II, or 25 uM of MEKinh until stage 28 was reached with replenishment every two hours. For activation of piezo1, embryos were incubated at stage 28 with 25 uM of Yoda 1 for ten minutes.

### Immunofluorescence Staining

*Xenopus tropicalis* embryos were fixed at stage 28 with either 4% paraformaldehyde for 20 minutes or 2% trichloroacetic acid for 10 minutes, followed by phosphate-buffered saline (PBS 1X) rinses and permeabilization with 0.2% PBST (PBS 1X and 0.2% Triton X-100) for 15-20 minutes at room temperature (RT). Embryos were then blocked in 5% bovine serum albumin (BSA) for 1 hour at RT, followed by incubation with the primary antibody(s) overnight at 4 degrees Celcius. After a couple of 0.2%PBST washes at RT, the embryos were then incubated with the corresponding secondary antibody(s) and phalloidin for 1 hour at RT, followed by nuclear staining with Hoechst for 10 minutes at RT and mounting onto a glass slide. We used the following primary antibodies: rabbit polyclonal anti-Piezo1 (Novus Biologicals, 1:50 dilution), Rabbit monoclonal anti-active Yap1 (Abcam, 1:200 dilution), rabbit monoclonal Erk ½ (cell signaling, 1:200 dilution), rabbit monoclonal phospho-Erk ½ (cell signaling, 1:200 dilution), mouse monoclonal anti-acetylated α-tubulin (sigma, 1:1000 dilution), mouse monoclonal anti-!-tubulin (sigma, 1:200 dilution).

### Western Blot and Analysis

Embryos were collected and lysed using a lysis buffer (100x protease inhibitor + 1x Tris-Buffered Saline (TBS), 1:100 dilution) and homogenized prior to centrifugation at 4 degrees Celcius for 15 minutes. Lysates were carefully collected from the supernatant, to which the sample buffer (4X Laemmli buffer + 50ul of beta-mercaptoethanol) was added to a final dilution of 1x, thoroughly mixed, and boiled at 95 degrees Celsius for ten minutes. The boiled samples were then stored at -20 degrees Celcius for future use or were allowed to cool prior to immunoblotting. The standard SDS-PAGE gel electrophoresis and western blot procedures were followed using the BioRad Trans-Blot 20 Mini Protean Tetra System, 4%-15% bis-acrylamide crosslinked gels, and polyvinylidene fluoride (PVDF) microporous membranes. After gel transfer, the membranes were probed with the respective primary antibody overnight at 4 degrees Celcius and the secondary antibody for 1-2 hours at RT with intermittently repeated washes with TBS with 0.1% Triton X-100 prior to chemiluminescent imaging using the gel imager (Azure Biosystems). The images were then analyzed using Image J/Fiji prior to assembly in Adobe Illustrator.

### Whole mount in-situ hybridization and HCR RNA-FISH

Whole mount in situ hybridization on stage 25 *X. tropicalis* embryos was performed as previously described (54). For HCR RNA-FISH experiments, the mRNA probes for *piezo1*, *foxj1,* and *myb* were synthesized by Molecular Instruments. After fixation at stage 25 with an aldehyde fixative (MEMFA) and dehydration with 100% ethanol, the embryos were rehydrated through multiple 5-minute washes with PBS-Tween (1X PBS+0.1% Tween) and permeabilized with Proteinase K for 10 minutes at RT, followed by charge neutralization with acetic anhydride in triethanolamine at RT. Embryos were re-fixed with 4% PFA in PBS-Tween (1x PBS + 0.1% Tween) for 20 minutes at RT. The embryos were then washed multiple times in PBS-Tween prior to pre-hybridization in the probe hybridization buffer (Molecular Instruments, CA) at 37C for 30 minutes, followed by hybridization of the samples with the mRNA probe (1:250 dilution in probe hybridization buffer) overnight at 37C. After multiple 15-minute washes with the wash buffer (Molecular Instruments, CA) at 37C, the samples were washed with 5X SSCT (5X SSC + 0.1% Tween) at RT and pre-amplified in the amplification buffer for 30 minutes at RT. For amplification, prepare the hairpin solution by snap-cooling the hairpins (Molecular Instruments, CA) followed by incubation of the samples in the hairpin solution (hairpins in amplification buffer, 1:50 dilution) overnight in the dark at RT. The excess hairpins are then removed with multiple washed using 5X SSCT. The samples were then processed for immunofluorescence staining and visualization.

### Data Acquisition and Microscopy

Post-staining embryos/animal explants were visualized using a Leica DMi8 SP8 confocal microscope using a 40x oil immersion objective (1.3 NA). For the acquisition of super-resolution images, the stained embryos were imaged using a Leica Stellaris 8 with FALCON and TauSTED microscope with a 100x oil immersion objective (1.4 NA). Images were captured at either 3x or 4x zoom and analyzed using Image J/Fiji prior to assembly in Adobe Illustrator.

### Data Analysis and Statistics

All experiments were performed at least three times, with measurements and analyses performed on at least 4 to 5 embryos per group. The IMARIS software and Fiji were used for centriole number quantification. Apical area measurements were acquired by marking the MCC boundary in Fiji with the polygon tool and measuring the area within the region of interest. Intensity measurements were recorded in Fiji with the help of either the rectangular tool in the case of western blots or the polygon tool for medial actin intensity measurements, as described in Kulkarni et al. (2018) (*20*). Statistical analyses were performed using GraphPad Prism version 10. The statistical analysis method, p-value, and significance are mentioned in the figure legends.

## ACKNOWLEDGEMENTS

We thank Dr. Karen Hirschi for sharing the Leica SP8 confocal microscope and the Advanced Microscope Facility at the University of Virginia. We also want to extend our heartfelt thanks to Douglas W. DeSimone for his valuable feedback on experiments. We also thank Dr. Sameer Bajikar, Dr. Douglas DeSimone, and Dr. Moe R. Mahjoub for reading and providing critical feedback on the manuscript.

## AUTHORS CONTRIBUTIONS

Conceptualization: SSK, VN Methodology: SSK, VN Investigation: VN, VGR, AA Visualization: VN Supervision: SSK Writing – original draft: SSK, VN

## DATA AND MATERIALS AVAILABILITY

All data and materials that support the findings from the study are available from the corresponding author upon request.

## SUPPLEMENTARY MATERIAL

**Supplemental Figure 1.**
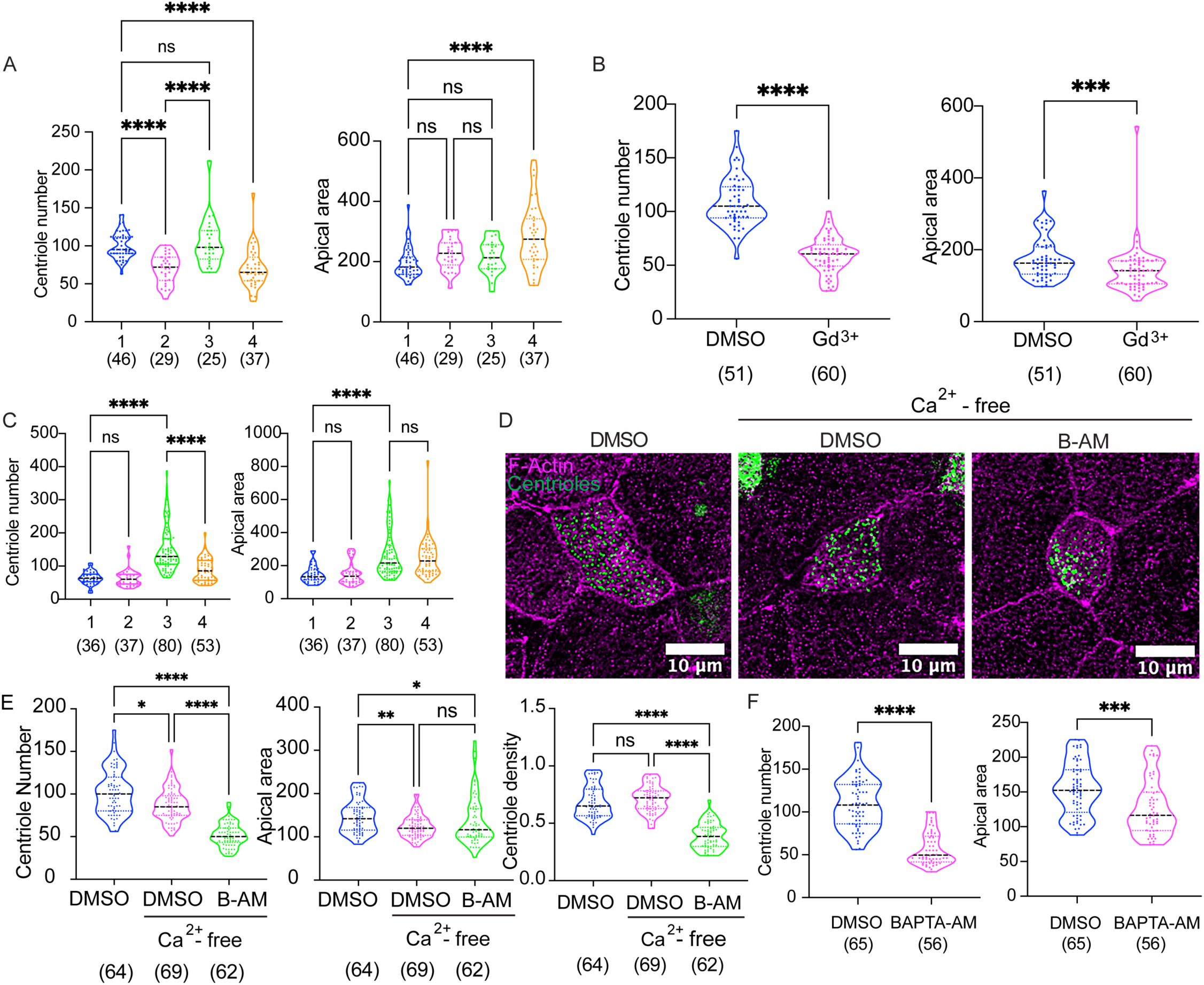
Role of Calcium in centriole amplification in *Xenopus* MCCs. A. Plot shows centriole number and apical area per MCC for all four cases. Kruskal-Wallis test followed by multiple comparisons was performed for statistical analysis between groups (p>0.05 (ns), ***p<0.001, and ****p<0.0001). Data shown is from three independent experiments; the number of MCCs from at least 5 embryos per trial is indicated in parentheses below each group. B. Plot depicts comparison of centriole number and apical area per MCC between control and Gd^3+^-treated stage 28 embryos. Mann-Whitney non-parametric t-test was used for statistical analysis (****p<0.0001). Data shown is from three independent experiments; the number of MCCs from at least 5 embryos per trial is indicated in parentheses below each group. C. Plot shows comparison of the centriole number and apical area per MCC between the different groups: 1, Untethered; 2, Untethered, Gd^3+^; 3, Tethered; 4, Tethered, Gd^3+^. Kruskal-Wallis test followed by multiple comparisons was performed for statistical analysis between groups (p>0.05 (ns), and ****p<0.0001). Data shown is from three independent experiments; the number of MCCs from at least 5 animal explants per experiment is indicated in parentheses below each group. D. Representative IF images of stage 28 MCCs that are either treated with a vehicle control (DMSO), treated with the vehicle control (DMSO, Ca^2+^-free) in the presence of calcium-free medium, or treated with BAPTA-AM in the presence of calcium-free medium (BAPTA-AM, Ca^2+^-free), expressing Chibby-BFP (centrioles, green) and are co-stained with Phalloidin (Magenta). Scale bar: 10um. E. Plot depicts a comparison of centriole number, apical area, and centriole density (centriole number/apical area) per MCC for all three groups in (D). Kruskal-Wallis test followed by multiple comparisons was performed for statistical analysis between groups (p>0.05 (ns), *p<0.05, **p<0.01, and ****p<0.0001). Data shown is from three independent experiments; the number of MCCs from at least 5 embryos per trial is indicated in parentheses below each group. F. Plot depicts a comparison of centriole number and apical area per MCC for DMSO-treated control (DMSO) and BAPTA-AM-treated (B-AM) stage 28 embryos. Mann-Whitney non-parametric t-test was used for statistical analysis (****p<0.0001). Data shown is from three independent experiments; the number of MCCs from at least 5 embryos per trial is indicated in parentheses below each group.

**Supplemental Figure 2.**
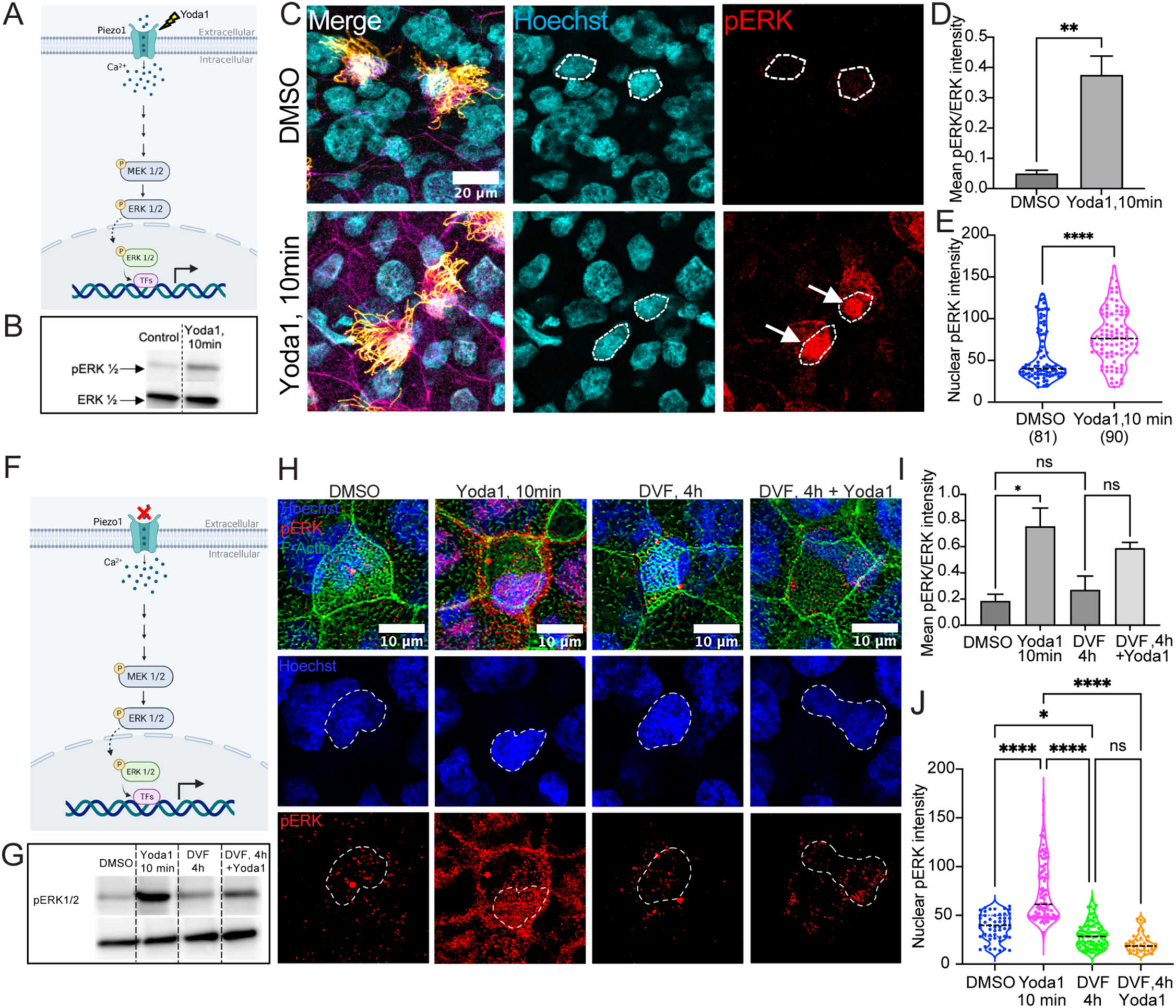
Piezo1-mediated calcium activates Erk1/2 in Xenopus MCCs. A. The schematic represents the mechanism of Piezo1 activation via Yoda1 and subsequent activation of the ERK ½ pathway, indicated by the nuclear translocation of phosphorylated ERK ½ to the nucleus. B. Western analysis showing the effect of Yoda1 (25 uM) treatment on ERK ½ phosphorylation. Embryos were treated with the vehicle control (DMSO) or Yoda1 for 10 minutes. The treated embryos were collected at stage 28 and compared for phospho-ERK ½ and total ERK ½ expression. The representative data shown is from one out of three independent experiments. C. Representative IF images show nuclear expression of phospho-ERK ½ (pERK, red) in MCCs across stage 28 embryos that were either treated with the vehicle control (DMSO) or Yoda1 for ten minutes (Yoda1, 10min). Embryos were co-stained with Phalloidin (F-Actin, magenta), acetylated tubulin (cilia, gold), and Hoechst (nuclei, cyan). Scale bar is 20 um. D. The plot shows the mean intensity of phospho-ERK ½ versus total ERK ½ across the different treatment groups in (B). Student t-test was performed for statistical analysis between groups (**p<0.01). Data is from three independent experiments; 30 embryos were used per group per trial. E. The plot shows the intensity of nuclear phospho-ERK ½ per MCC in stage 28 embryos that were either DMSO-treated (DMSO) or treated with Yoda1 for ten minutes (Yoda1, 10min). Mann-Whitney test was performed for statistical analysis between groups (****p<0.0001). Data shown is from three independent experiments. The number of MCCs used for each group from every trial is indicated in parentheses. At least 5 embryos were used per group in every experiment. F. Schematic represents the mechanism by which the extra-cellular calcium is removed using a divalent-free medium (DVF). G. Western analysis shows the effect of Yoda1 (25 uM) treatment in the absence of extracellular calcium on ERK ½ phosphorylation. Stage 28 embryos were either treated with the vehicle control (DMSO), Yoda1 for 10 minutes (Yoda1, 10min), the divalent-free medium for 4 hours (DVF, 4h), or pre-treated with the divalent-free medium for 4 hours followed by Yoda1-treatment for 10 minutes in divalent-free medium (DVF, 4h, + Yoda1). The treated embryos were collected at stage 28 and compared for phospho-ERK ½ and total ERK ½ expression. The representative data shown is from one out of three independent experiments. H. Representative IF images show nuclear expression of phospho-ERK ½ (pERK, red) in MCCs across different groups in (G). Embryos were co-stained with Phalloidin (F-Actin, green), and Hoechst (nuclei, cyan). Scale bar is 20 um. I. Plot shows mean intensity of phospho-ERK ½ versus total ERK ½ across the different treatment groups in (G). One-Way ANOVA test followed by multiple comparisons was performed for statistical analysis between groups (p>0.05 (ns), **p<0.01, and ****p<0.0001). Data shown is from four independent experiments; 30 embryos were used per group per trial. J. Plot shows intensity of nuclear phospho-ERK ½ per MCC in stage 28 embryos across different groups in (G). Kruskal-Wallis test followed by multiple comparisons was performed for statistical analysis between groups (****p<0.0001). Data shown is from three independent experiments. The number of MCCs used for each group from every trial is indicated in parentheses. At least 5 embryos were used per group in every experiment.

**Supplemental Figure 3.**
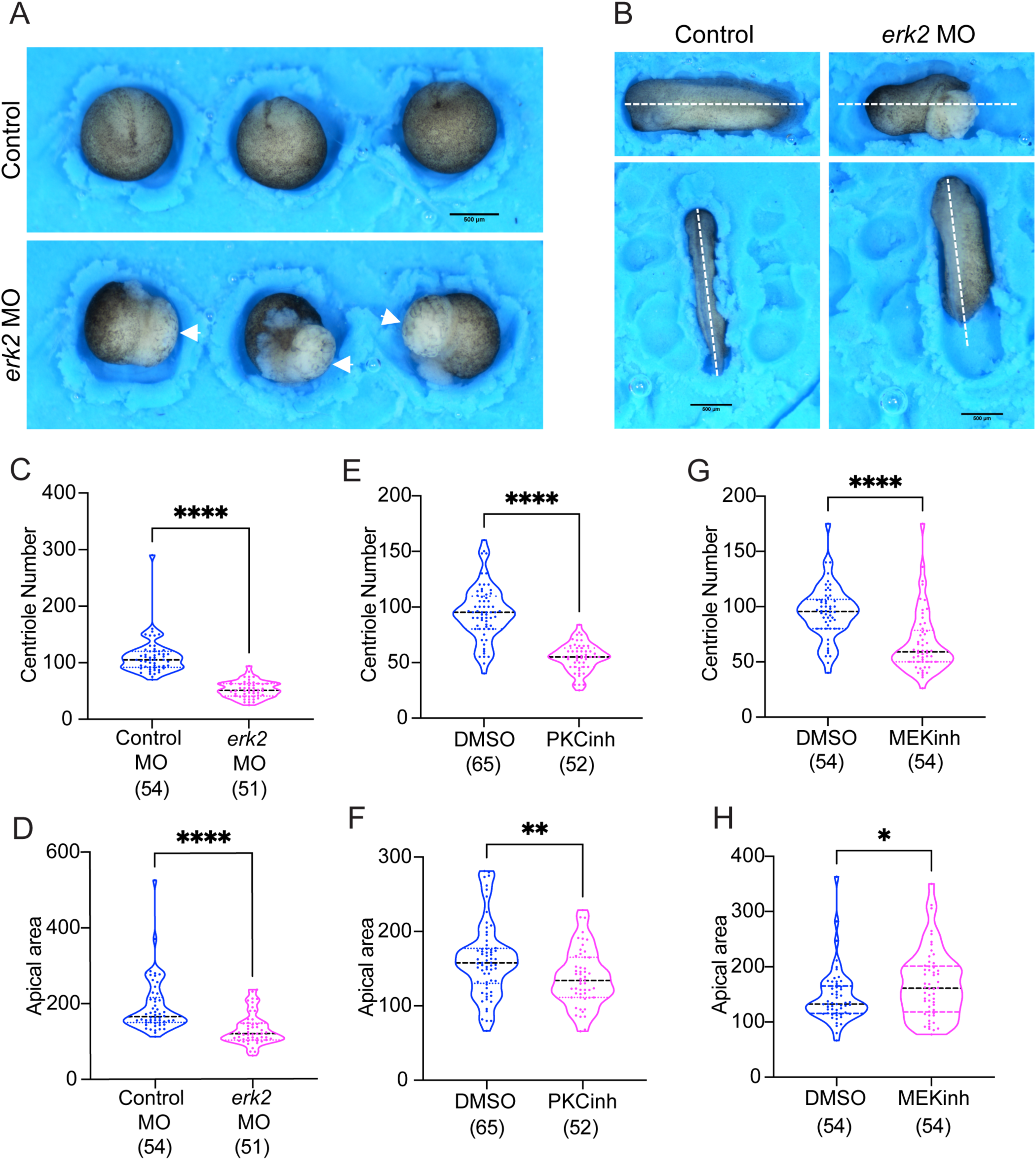
Erk1/2 is essential for embryonic development and MCC morphogenesis in *Xenopus*. A. Comparison between stage 16 *erk2*-depleted (*erk2* MO) and uninjected embryos. Representative images indicate gastrulation defects (open blastopore; white arrows) with knockdown of *erk2* via *erk2* MO. Scale bar: 500um B. Comparison between stage 28 *erk2*-depleted (*erk2* MO) and uninjected embryos. Representative images indicate anterior-posterior (AP) defects (shortened AP axis; dotted lines) with knockdown of *erk2* via *erk2* MO. Scale bar: 500um C. Plot depicts a comparison of centriole number per MCC between standard control (Control MO) and *erk2*- depleted (*erk2* MO) stage 28 embryos. Mann-Whitney non-parametric t-test was used for statistical analysis (****p<0.0001). Data shown is from three independent experiments; the number of MCCs from at least 5 embryos per trial is indicated in parentheses below each group. D. Plot depicts comparison of apical area per MCC between standard control (Control MO) and *erk2* KD via *erk2* MO (*erk2* MO) stage 28 embryos. Mann-Whitney non-parametric t-test was used for statistical analysis (****p<0.0001). Data shown is from three independent experiments; the number of MCCs from at least 5 embryos per trial is indicated in parentheses below each group. E. Plot depicts a comparison of centriole number per MCC between vehicle control (DMSO) and PKC inhibitor-treated (PKCinh) stage 28 embryos. Mann-Whitney non-parametric t-test was used for statistical analysis (****p<0.0001). Data shown is from three independent experiments; the number of MCCs from at least 5 embryos per trial is indicated in parentheses below each group. F. Plot depicts a comparison of apical area per MCC between vehicle control and PKC inhibitor-treated (PKCinh) stage 28 embryos. Mann-Whitney non-parametric t-test was used for statistical analysis (****p<0.0001). Data shown is from three independent experiments; the number of MCCs from at least 5 embryos per trial is indicated in parentheses below each group. G. Plot depicts comparison of centriole number per MCC between vehicle control (DMSO) and MEK ½ inhibitor-treated (MEKinh) stage 28 embryos. Mann-Whitney non-parametric t-test was used for statistical analysis (****p<0.0001). Data shown is from three independent experiments; the number of MCCs from at least 5 embryos per trial is indicated in parentheses below each group. H. Plot depicts comparison of apical per MCC between control and MEK ½ inhibitor-treated (MEKinh) stage 28 embryos. Mann-Whitney non-parametric t-test was used for statistical analysis (*p<0.01). Data shown is from three independent experiments; the number of MCCs from at least 5 embryos per trial is indicated in parentheses below each group.

**Supplemental Figure 4.**
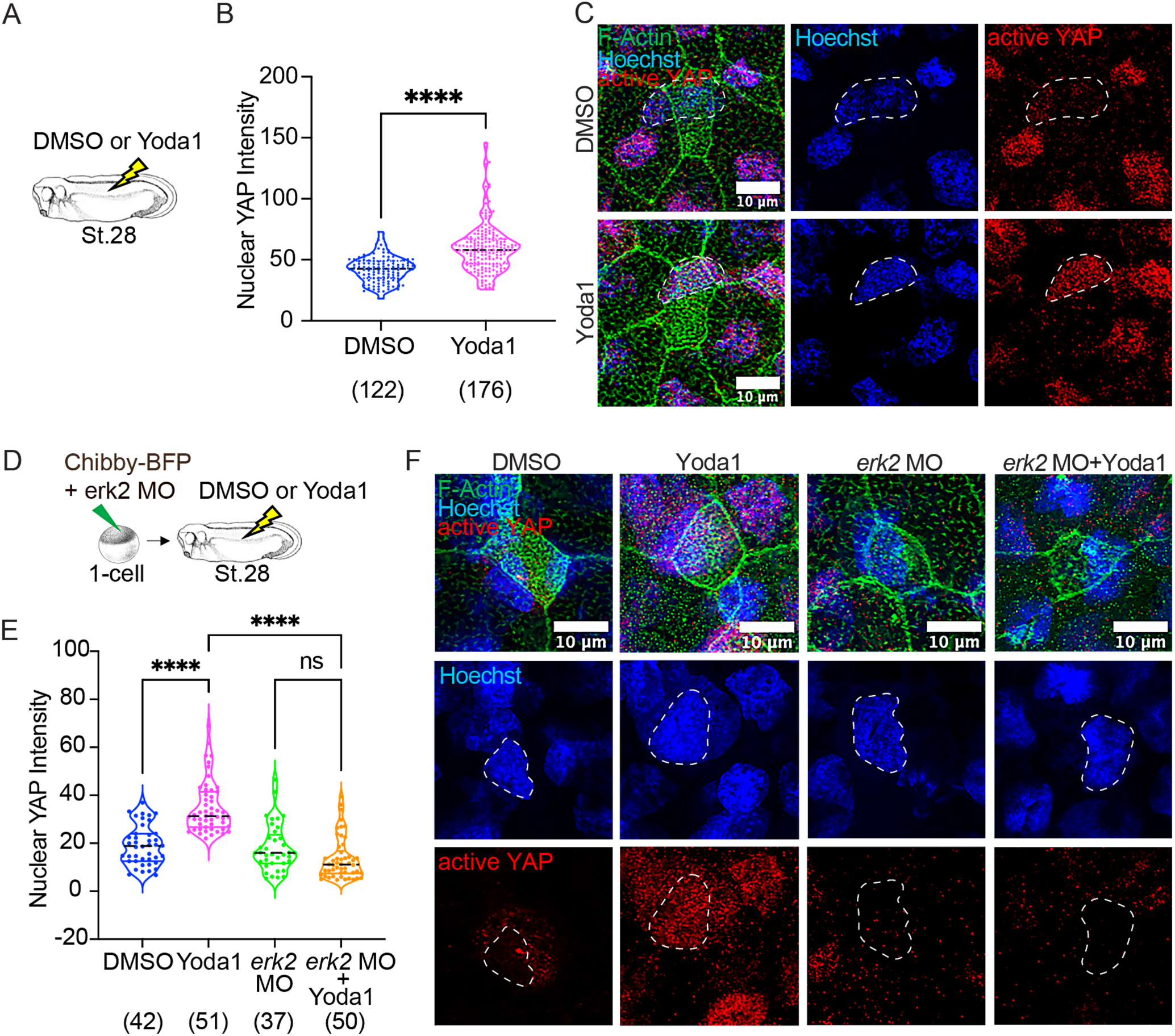
Piezo1 regulates Yap1 activity via Erk2 in *Xenopus* MCCs. A. Schematic represents the experimental set-up involving treatment of stage 28 embryos with either the vehicle control, DMSO or Piezo1 activator, Yoda1. B. Plot shows nuclear YAP intensity with Piezo1 activation via Yoda1. Stage 28 embryos were either treated with the vehicle control (DMSO) or Yoda1 (25uM) for ten minutes. The nuclear YAP intensity values were recorded with the help of the polygon tool and intensity measurement features in Fiji for further analysis. C. Representative IF images show the effect of Piezo1 activation on the nuclear translocation of YAP in MCCs. Stage 28 embryos were either treated with the vehicle control (DMSO) or Yoda1 (25uM) for ten minutes and stained for active-YAP (red) and phalloidin (green). Nuclei were counterstained with Hoechst (blue). Scale bar is 10 um. D. Schematic represents the experimental set-up for the effect of Piezo1 activation on nuclear YAP localization in *erk2* KD embryos. Embryos were injected with the *erk2* MO and centriole marker (Chibby- BFP, 100 pg) at 1-cell stage. Stage 28 embryos were then either treated with the vehicle control (DMSO) or Yoda1 (25 uM) for ten minutes and processed for IF staining and visualization. E. Plot shows intensity of nuclear YAP1 (active-YAP) per MCC across the different treatment groups. Embryos that were injected with *erk2* MO (20ng) at the 1-cell stage are either treated with Yoda1 for 10 minutes (erk2 MO+Yoda1) or left untreated (erk2 MO) and compared against the unperturbed embryos treated with the vehicle control (DMSO) and those treated with Yoda1 for 10 minutes (Yoda1, 10min). Kruskal-Wallis test followed by multiple comparisons was performed for statistical analysis between groups (p>0.05 (ns), and ****p<0.0001). Data shown is from three independent experiments; the number of MCCs from at least 5 embryos per trial is indicated in parentheses below each group. F. Representative IF images show the effect of Piezo1 activation via Yoda1 (25 uM) on the nuclear localization of YAP in *erk2* KD MCCs from stage 28 embryos. Embryos that were injected with *erk2* MO (20ng) at the 1-cell stage are either treated with Yoda1 for 10 minutes (erk2 MO+Yoda1) or left untreated (erk2 MO) and compared against the unperturbed embryos treated with the vehicle control (DMSO) and those treated with Yoda1 for 10 minutes (Yoda1, 10min). Embryos are stained for active-YAP (red) and co-stained with Phalloidin (F-Actin, green) and Hoechst (nuclei, blue). Scale bar is 10um.

**Supplemental Figure 5.**
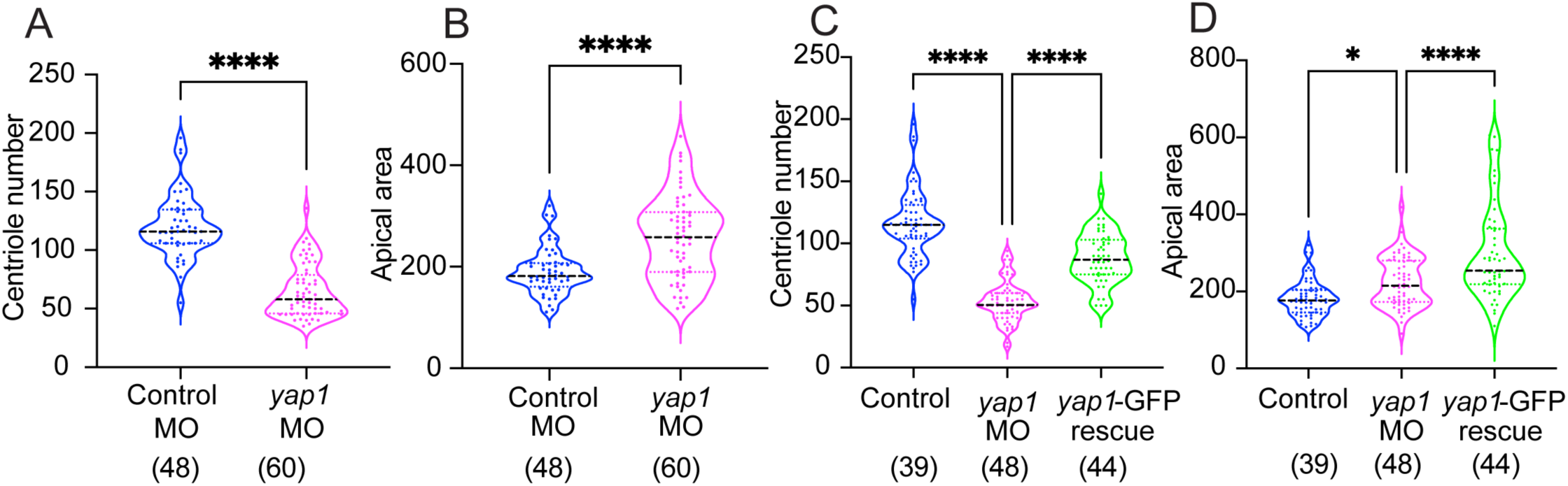
Yap control of centriole amplification in *Xenopus* MCCs. A. Plot depicts comparison of centriole number per MCC between standard control (Control MO) and *yap1* KD (*yap1* MO) stage 28 embryos. Mann-Whitney non-parametric t-test was used for statistical analysis (****p<0.0001). Data shown is from three independent experiments; the number of MCCs from at least 5 embryos per trial is indicated in parentheses below each group. B. Plot depicts comparison of apical area per MCC between standard control (Control MO) and *yap1* KD (*yap1* MO) stage 28 embryos. Mann-Whitney non-parametric t-test was used for statistical analysis (****p<0.0001). Data shown is from three independent experiments; the number of MCCs from at least 5 embryos per trial is indicated in parentheses below each group. C. Plot depicts comparison of centriole number per MCC between unperturbed controls, yap1 KD (*yap1* MO), and *yap1*-rescued stage 28 MCCs (yap1-GFP rescue). Kruskal-Wallis test followed by multiple comparisons was performed for statistical analysis between groups (****p<0.0001). Data shown is from three independent experiments; the number of MCCs from at least 5 embryos per trial is indicated in parentheses below each group. D. Plot depicts comparison of apical area per MCC between unperturbed controls, yap1 KD (*yap1* MO), and *yap1*-rescued stage 28 MCCs (yap1-GFP rescue). Kruskal-Wallis test followed by multiple comparisons was performed for statistical analysis between groups (*p<0.05, and ****p<0.0001). Data shown is from three independent experiments; the number of MCCs from at least 5 embryos per trial is indicated in parentheses below each group.

**Supplemental Figure 6.**
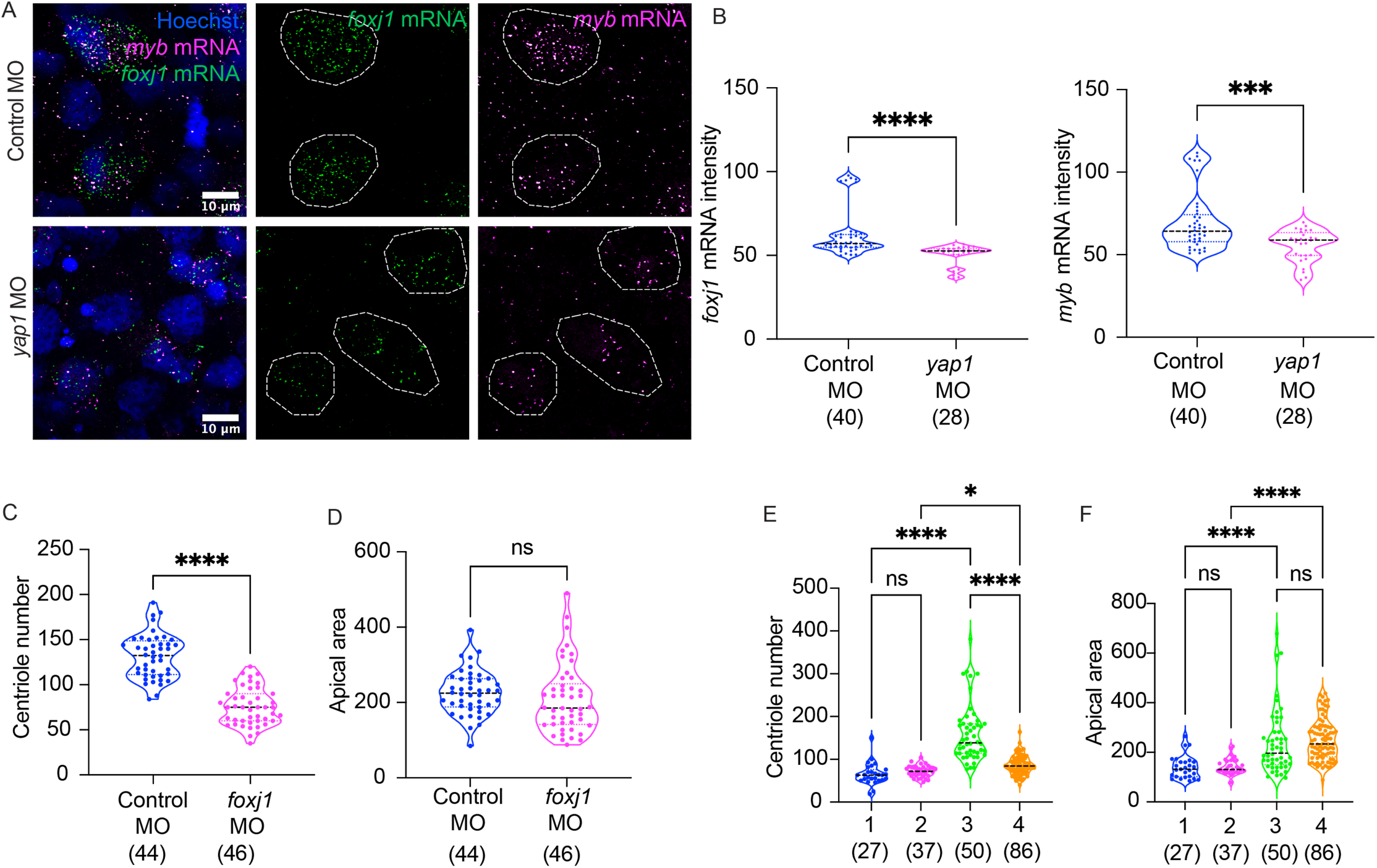
Yap-Foxj1 axis controls centriole amplification in *Xenopus* MCCs. A. Representative HCR RNA-FISH images of stage 25 MCCs that are either injected at 1-cell stage with the standard control (Control MO) or *yap1* morpholino (*yap1* MO). Embryos are collected at stage 25 for HCR RNA-FISH and visualization. Embryos are probed with Myb (*myb* mRNA, magenta) and Foxj1 (*foxj1* mRNA, green) HCR RNA probe. Nuclei are counterstained with Hoechst (blue). Scale bar: 10um. B. Plots show a comparison between control (Control MO) and yap1 KD (*yap1* MO) stage 25 MCCs for *myb* mRNA intensity (*right*), and *foxj1* mRNA intensity (*left*) per MCC. Mann-Whitney non-parametric t-test was used for statistical analysis (****p<0.0001). Data shown is from three independent experiments; the number of MCCs from at least 3 embryos per group per trial is indicated in the parentheses below each group. C. Plot depicts comparison of centriole number per MCC between control and foxj1 KD (*foxj1* MO) stage 28 embryos. Mann-Whitney non-parametric t-test was used for statistical analysis (****p<0.0001). Data shown is from three independent experiments; the number of MCCs from at least 5 embryos per trial is indicated in parentheses below each group. D. Plot depicts comparison of apical area per MCC between control and foxj1 KD (*foxj1* MO) stage 28 embryos. Mann-Whitney non-parametric t-test was used for statistical analysis (p>0.05 (ns)). Data shown is from three independent experiments; the number of MCCs from at least 5 embryos per trial is indicated in parentheses below each group. E. The Plot compares the centriole number per MCC between stage 28 animal explants with foxj1 depletion (via *foxj1* MO at 1-cell stage). The tethered explant depleted of foxj1 (4: Tethered, *foxj1* MO) is compared against the uninjected tethered control (3: Tethered) and the untethered groups (Untethered (1) and Untethered, *foxj1* MO (2)). Kruskal-Wallis test followed by multiple comparisons was performed for statistical analysis between groups (p>0.05 (ns), and ****p<0.001). Data shown is from three independent experiments; the number of MCCs from at least 5 embryos per trial is indicated in parentheses below each group. F. The Plot compares the apical area per MCC between the different groups in (F). Kruskal-Wallis test followed by multiple comparisons was performed for statistical analysis between groups (p>0.05 (ns), and ****p<0.001). Data shown is from three independent experiments; the number of MCCs from at least 5 embryos per trial is indicated in parentheses below each group.

**Supplemental Figure 7.**
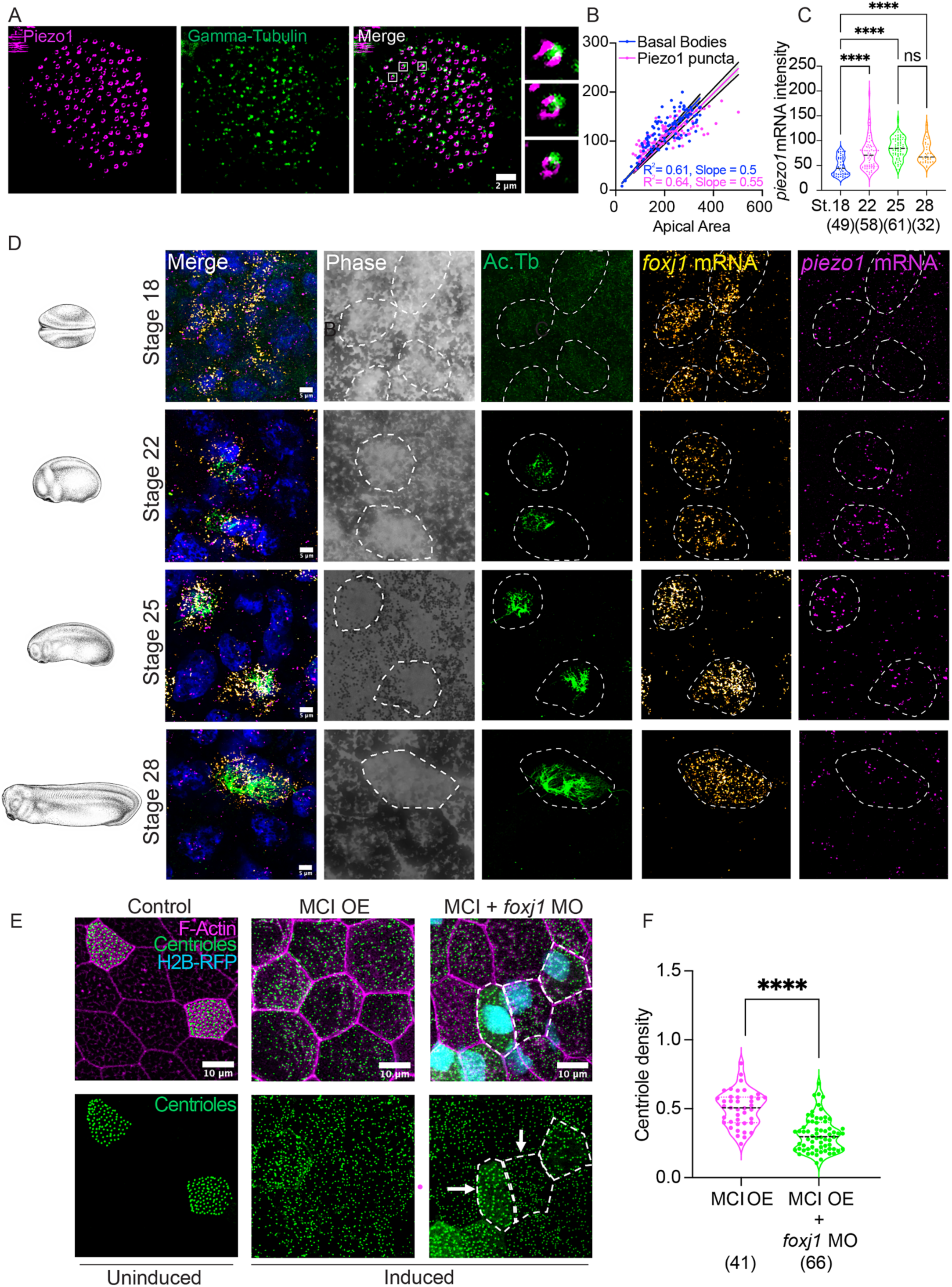
Piezo1 and centrioles are regulated MCIDAS-Foxj1 in *Xenopus* MCCs. A. Representative image of a mature multiciliated cell (MCC) from a stage 28 embryo, immuno-stained with Piezo1 antibody (magenta) and gamma-tubulin (green). Scale bars: 2 um. B. Scatterplot depicting the correlation between piezo1 puncta (magenta dots) and centrioles (blue dots) in MCCs with increase in apical area across the different developmental stages, namely, stage 18, 22, 25, and 28. C. Plot shows piezo1 mRNA intensity across the different embryonic developmental stages: 18, 22, 25, and 28, depicting the mRNA expression levels of piezo1 using HCR-RNA FISH. Data shown is from three independent experiments; Kruskal-Wallis test followed by multiple comparisons was performed for statistical analysis between groups (ns: p>0.05, and ****p<0.0001). Data shown is from three independent experiments; the number of MCCs from at least 4 embryos per trial is indicated in parentheses below each group. D. Representative IF images from HCR-RNA FISH experiments for embryos that were hybridized with the piezo1 mRNA probe and assessed for piezo1 mRNA expression levels (magenta) across the different stages (18, 22, 25, and 28). The foxj1 mRNA probe was used to trace MCCs (gold). The embryos were co-stained with acetylated tubulin (green), and the transmission phase channel was obtained to detect cell boundaries (grey). Scale bar: 5um. E. Representative IF images of stage 28 MCCs, expressing the centriole marker (Chibby-BFP, green), are either uninduced (Control), induced for multicilin over-expression (*MCI* OE), or induced for multicilin over- expression and depleted of foxj1 (*MCI* + *foxj1* MO) and co-stained with Phalloidin (magenta). Embryos were injected with multicilin (*MCI*, 50 pg) and centriole marker (Chibby-BFP, 100 pg) at 1-cell stage, followed by mosaic depletion of foxj1 via *foxj1* MO along with a nuclear tracer (H2B-RFP, 100 pg) in 1 of the 4 cells at the 4-cell stage. Embryos were then induced for multicilin over-expression using dexamethasone (0.1 mg/ml) at stage 10 and collected at stage 28 for immunofluorescence and visualization. MCCs depleted of foxj1 are traced by their expression of the nuclear tracer (H2B-RFP, cyan), as indicated by the arrows in the images. Scale bar: 10um. F. The plot shows comparison of the centriole density per MCC between multicilin over-expressed MCCs (*MCI* OE) and multicilin over-expressed MCCs that are depleted of foxj1 (*MCI* + *foxj1* MO). Mann-Whitney non-parametric t-test was used for statistical analysis (****p<0.0001). Data shown is from three independent experiments; the number of MCCs from at least 5 embryos per trial is indicated in parentheses below each group.

